# Deep mining of early antibody response in COVID-19 patients yields potent neutralisers and reveals high level of convergence

**DOI:** 10.1101/2020.12.29.424711

**Authors:** Georgia Bullen, Jacob D. Galson, Pedro Villar, Lien Moreels, Line Ledsgaard, Giada Mattiuzzo, Gareth Hall, Emma M. Bentley, Edward W. Masters, David Tang, Sophie Millett, Danielle Tongue, Richard Brown, Ioannis Diamantopoulos, Kothai Parthiban, Claire Tebbutt, Rachael Leah, Krishna Chaitanya, Deividas Pazeraitis, Sachin B. Surade, Omodele Ashiru, Lucia Crippa, Richard Cowan, Matthew W. Bowler, Jamie I. Campbell, Wing-Yiu Jason Lee, Mark D. Carr, David Matthews, Paul Pfeffer, Simon E. Hufton, Kovilen Sawmynaden, Jane Osbourn, John McCafferty, Aneesh Karatt-Vellatt

## Abstract

Passive immunisation using monoclonal antibodies will play a vital role in the fight against COVID-19. Until now, the majority of anti-SARS-CoV-2 antibody discovery efforts have relied on screening B cells of patients in the convalescent phase. Here, we describe deep-mining of the antibody repertoires of hospitalised COVID-19 patients using a combination of phage display technology and B cell receptor (BCR) repertoire sequencing to isolate neutralising antibodies and gain insights into the early antibody response. This comprehensive discovery approach has yielded potent neutralising antibodies with distinct mechanisms of action, including the identification of a novel non-ACE2 receptor blocking antibody that is not expected to be affected by any of the major viral variants reported. The study highlighted the presence of potent neutralising antibodies with near germline sequences within both the IgG and IgM pools at early stages of infection. Furthermore, we highlight a highly convergent antibody response with the same sequences occurring both within this study group and also within the responses described in previously published anti-SARS-CoV-2 studies.

## Introduction

In the past two decades, three major virus outbreaks caused by coronaviruses have emerged. The latest, COVID-19 caused by Severe Acute Respiratory Syndrome Coronavirus 2 (SARS-CoV-2), has resulted in a pandemic with over 78 million people infected and causing over 1.7 million deaths. The majority of drug development efforts have been focused on vaccine development, with 51 and 163 programs going through clinical and preclinical evaluation respectively at the beginning of December 2020 (Source: WHO). Despite the accelerated development timeframes of the leading vaccine candidates, the world is still one or two years away from attaining population immunity due to the manufacturing and logistical challenges of mass vaccinating billions of people. Therefore, monoclonal antibodies can become a key component in the early fight against COVID-19. Viral neutralising antibodies can offer a two-in-one approach; they can be used both to treat symptomatic individuals following acute exposure, and as a prophylactic to protect healthcare workers and at-risk groups, including individuals who respond poorly to vaccines.

There are 13 experimental anti-SARS-CoV-2 monoclonal antibody treatments undergoing clinical trials with bamlanivimab and a cocktail containing casirivimab and imdevimab receiving Emergency Use Approval from the Food and Drug Administration (FDA). Most well-characterized and highly potent neutralising antibodies in both clinical and preclinical development target the SARS-CoV-2 S protein and its receptor binding domain (RBD) *(1)*. The majority of these neutralising antibodies have been derived from single cell screening of memory B cells from COVID-19 patients in their convalescent phase of disease (blood samples were collected on average 32 days after the onset of symptoms, Table S1).

Here, we present a complementary discovery approach (Fig. 1), which combined phage display technology and BCR repertoire sequence analysis. This approach allowed the isolation of hundreds of anti-SARS-CoV-2 antibodies from the antibody repertoires of patients in the acute phase of disease (blood samples collected on average 11 days (range 4-20) after the onset of symptoms) and also to study the early antibody response. A comprehensive search of patient-derived phage display libraries combined with high-throughput biochemical and functional screening resulted in the discovery of highly potent neutralising antibodies, with diverse epitopes and distinct mechanisms of action. An emphasis on developability testing as part of early discovery screening facilitated selection of a panel of well characterised antibodies with biophysical properties de-risked for downstream development and manufacturing. Armed with the knowledge of functional binding activity, antibody sequences discovered by phage display were co-clustered with whole BCR repertoire sequencing from the patients and published antibodies to understand the nature and dynamics of the early antibody response, including the isotype usage, clonal expansion and the level of convergence.

**Fig. 1.**
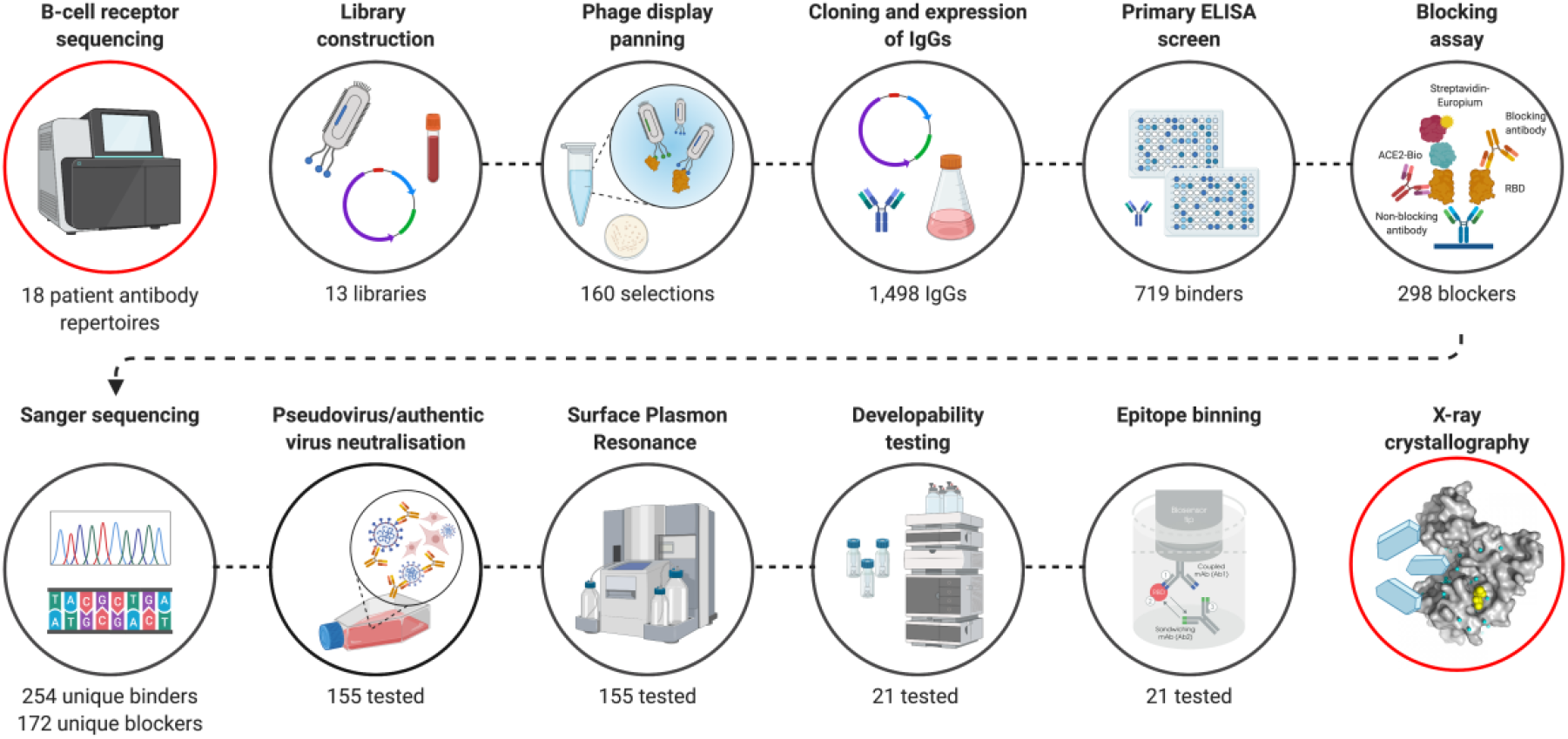
Overview of SARS-CoV-2 antibody discovery and analysis of patient response to COVID-19. Antibody genes isolated from the PBMC’s of 18 COVID-19 patients were used to for B cell receptor repertoire sequencing and to construct phage display libraries. Characterization of phage display derived antibodies using high throughput expression, primary binding assay and biochemical ACE-2 receptor blocking assays and DNA sequencing resulted in testing of 155 unique antibodies for pseudovirus neutralization and surface plasmon resonance. A final panel of 21 antibodies were subjected to authentic virus neutralization, epitope binding and developability assessment. Finally, structures of a complementary pair of antibodies with two different mechanism of viral neutralisation in complex with RBD were determined using X-ray crystallography. The V_H_ sequences of RBD binding and pseudoviral neutralising antibodies were co-clustered with whole BCR repertoire sequencing from the patients and published antibodies to understand the nature and dynamics of the early antibody response. Figure was prepared using BioRender.

## Results

### Isolation of anti-SARS-CoV-2 antibodies from patient derived libraries using phage display technology

Peripheral blood samples of 18 patients admitted to hospital with acute COVID-19 pneumonia were collected after obtaining informed consent. Patient demographics and clinical information relevant to their admission were also collected (Table S2). The patients experienced an average of 11 days (range 4-20 days) of symptoms prior to the day on which the blood sample was taken.

Initial B cell receptor (BCR) sequence analysis of the V_H_ repertoire of these patients revealed a strong convergent sequence signature. In order to link sequence data to information on binding properties, V_H_ populations from these donors were incorporated into phage display libraries in the form of single chain variable fragment (scFv) binding. By one approach we constructed hybrid libraries (of size 0.9 × 10^9^) where the variable heavy (V_H_) genes from the patient IgG repertoire were combined with a pre-existing library of variable light (V_L_) genes derived from healthy donors *(2)*. The second strategy focussed on creating fully patient-derived libraries (of size 1.5 × 10^9^) by assembling V_H_s and V_L_s on a donor-by-donor basis. Phage particles rescued from these libraries were used to carry out 3 rounds of panning on monomeric RBD and S1 ectodomains. The stringency of panning in each round was increased by reducing the antigen concentration to enrich for higher affinity binders.

Following three rounds of panning, the selected populations from both sets of libraries were sub-cloned *en masse* into an IgG expression vector while maintaining V_H_-V_L_ pairing from the selected scFvs. A total of 1498 clones were picked and expressed as IgG1 in Expi293F cells and screened using an affinity capture assay which eliminates the effect of expression variation between clones *(3, 4)*. Of the 1498 clones expressed, 589 antibodies originated from the phage panning against S1 and were screened against both RBD and S1, while the remaining 909 antibody clones from RBD panning were screened only on RBD. This screen yielded a total of 719 binders (48%). Surprisingly, a vast majority (87.3%) of the binders originating from S1 selections were also directed at RBD epitopes, resulting in a mere 46 non-RBD S1 binders. The preferential enrichment of antibodies to RBD over other S1 epitopes indicates that RBD is the key immunogenic region within the S1 protein. Although the further characterisation of the S1 binders led to identification of neutralising antibodies, these antibodies will be not discussed in detail in this manuscript.

### Functional, biochemical and biophysical characterisation of anti-SARS-CoV-2 antibodies

It is reported that the primary mechanism of the most potent SARS-CoV-2 neutralising antibodies is by blocking the RBD interaction with ACE2 cell surface receptor *(1, 5, 6)*. Therefore, we tested 394 RBD binders for their ability to block ACE2 in a biochemical assay (Fig. 2A), while carrying out the Sanger sequencing of these clones in parallel. Of these, 254 antibodies had a unique V_H_ and V_L_ CDR3 sequence and 172 unique antibodies showed >30% blocking in the biochemical assay. From this category, 121 unique antibody blockers were selected for viral neutralisation assays. In addition, 34 clones which failed to block the RBD-ACE2 interaction were progressed in the search for alternative mechanisms of action. This panel of 155 antibodies was subsequently purified from culture supernatants and screened in a lentiviral based SARS-CoV-2 pseudovirus assay at a concentration of 25 nM (3.5 μg/ml). 114 out of 121 (94.2%) ACE2 blocking antibodies showed >30% neutralisation. Among this group, 97 (80.2%) antibodies showed neutralisation activity exceeding 80%. Interestingly, a significant proportion of RBD-binding antibodies which failed to show ACE2 blocking in the biochemical assay (50%) showed neutralisation of pseudovirus, albeit significantly weaker than the ACE2 blockers (with 9/34 antibodies showing >50% neutralisation and only 1 antibody exceeding 80% neutralisation). Selected antibodies from the two different library approaches (“hybrid” versus fully patient derived) performed equally well and produced a similar number of blockers and neutralisers (Fig. 2B).

**Fig. 2.**
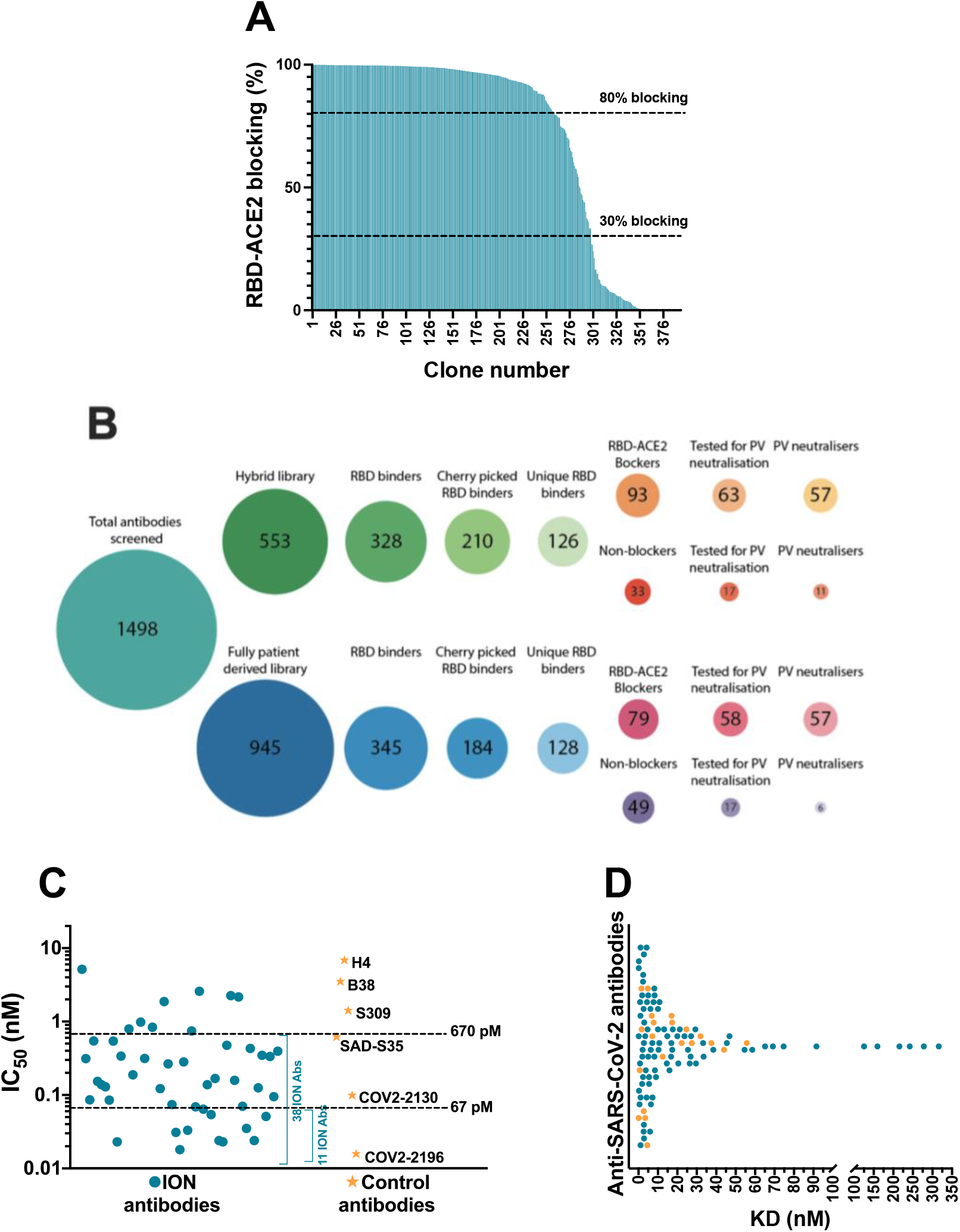
Biochemical and functional characterisation of SARS-CoV-2 antibodies. A) Screening of SARS-CoV-2 binders for RBD-ACE2 blocking activity. Antibodies were tested and ranked in a biochemical blocking assay for their ability to block the RBD-ACE2 interaction. B) Antibody screening process presented as bubble plots. Areas of bubbles are scaled relative to each other based on the number of antibodies that represent each bubble. C) Pseudovirus neutralising activity of 52 SARS-CoV-2 antibodies and control antibodies. D) Dot plot representing the 1:1 RBD binding affinities of SARS-CoV-2 antibodies measured using SPR. The antibodies that are part of the final panel of 21 are represented in orange dots.

The half maximal inhibitory concentrations (IC_50_) of the top 52 antibodies were determined (Fig. 2C and Fig. S1). This experiment also included a number of recently published SARS-CoV-2 neutralising antibodies including the potent antibody pair (COV2-2196 and COV2-2130) identified by the Vanderbilt University-AstraZeneca team *(7)*. Of the 52 antibodies tested, 38 were categorised as strong neutralisers with IC_50_ values below 670 pM (100 ng/ml). Amongst the control antibodies, COV2-2196 and COV2-2130 demonstrated best neutralisation with IC_50_ titres of 16 pM and 99 pM respectively. 11 of the most potent antibodies neutralised the pseudovirus with IC_50_ ranging from 18-67 pM (2.7 ng/ml to 10 ng/ml) hence matching or exceeding the best antibodies reported *(5, 7–11)*.

In parallel to the pseudovirus neutralisation assay, the binding kinetics of 155 antibodies were determined by high-throughput surface plasmon resonance (SPR). The affinities of these antibodies ranged from 70 pM to 316 nM, with most antibodies clustered in the range of 1-30 nM (Fig. 2D, Table S3). Comparison of the 52 clones tested for both affinity measurement and IC_50_ determination in pseudoviral neutralisation assay, showed poor correlation between affinity and neutralisation potency (Fig. S2A). In addition, the cross-reactivity of these antibodies to SARS-CoV-1 and MERS-CoV were also evaluated using a binding assay based on time resolved fluorescence (TRF). Only the antibodies that bound to SARS-CoV-1 or MERS-CoV with binding signals within 5-fold of SARS-CoV-2 were considered as cross-reactives. 29/155 (18.7 %) SARS-CoV-2 antibodies tested for cross-reactivity showed binding to SARS-CoV-1 RBD, which included 16 pseudovirus neutralisers (Fig. 3A). Unsurprisingly, none of the antibodies recognised MERS-CoV S1 which has very low sequence homology with SARS-CoV-2 (~20%). Interestingly, a significantly higher proportion of non-ACE2 blocking antibodies showed cross reactivity to SARS-CoV-1 (55.9%) than the ACE2 blockers (8.3%).

**Fig. 3.**
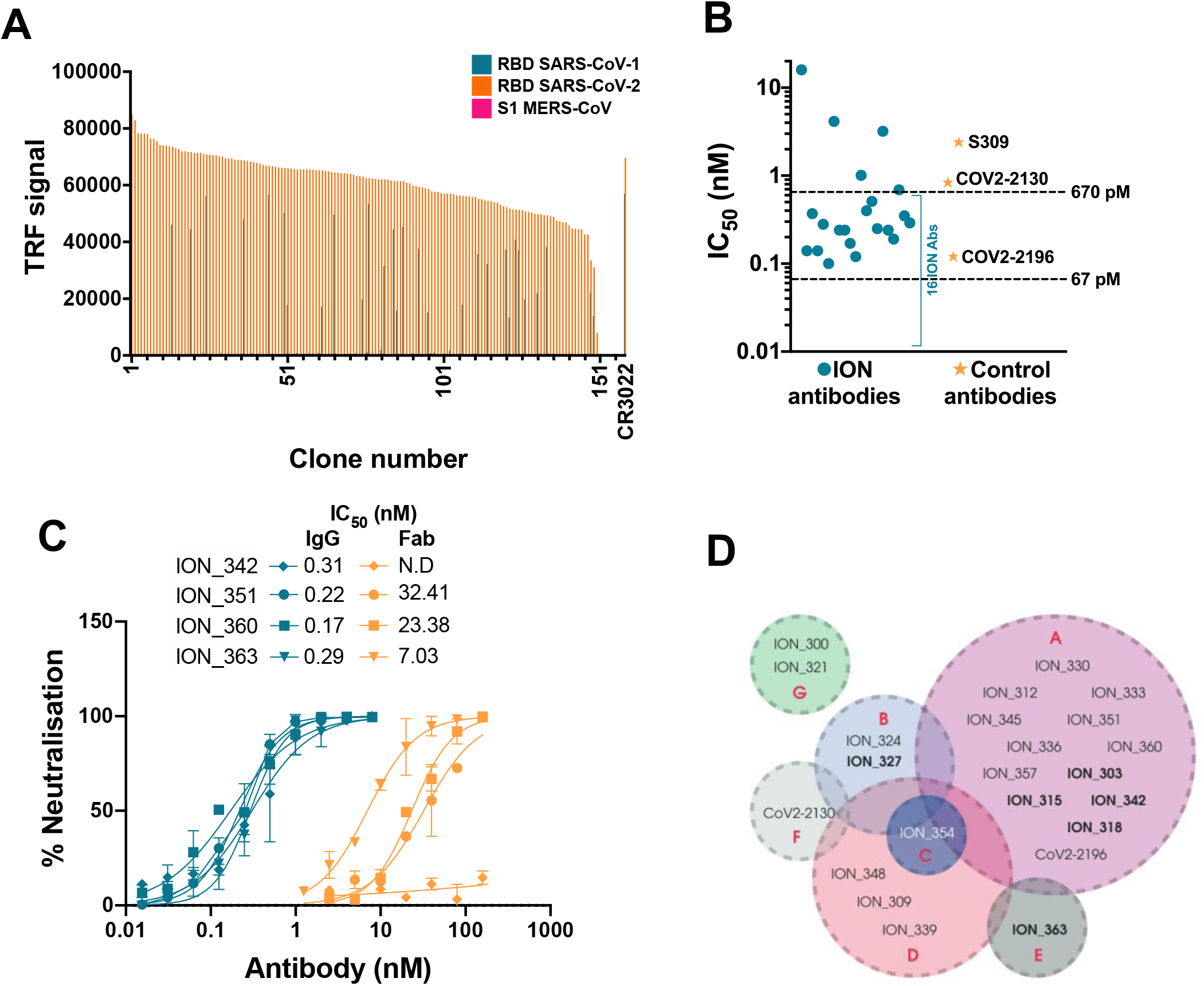
Detailed characterisation of SARS-CoV-2 antibodies. A) Evaluating the cross reactivity of 155 SARS-CoV-2 antibodies to SARS-CoV-1 RBD and MERS-CoV S1. CR3022, a previously published binding SARS-CoV-1 and SARS-CoV-2 cross reactive antibody was used as control. B) Authentic SARS-CoV-2 neutralising activity of a panel of 21 SARS-CoV-2 antibodies and control antibodies. C) Comparing authentic SARS-CoV-2 neutralising activity of 4 antibodies, in Fab and IgG format. D) Epitope binning of SARS-CoV-2 antibody panel using Octet BioLayer interferometry. The bins are labelled A to G. Antibodies in bold text denote capture molecules that exhibited unidirectionality.

Based on the potency of pseudovirus neutralisation, a final panel of 21 antibodies were selected for further characterisation. Given the abundance of potent neutralisers, any antibodies with poor transient expression and sequence liabilities (deamidation motifs in CDRs, unpaired cysteines and glycosylation motifs in V regions) were excluded at this stage. This panel also included two non-blocking neutralisers, which despite showing only moderate neutralisation in the pseudovirus assay were included due to their distinct mechanism of neutralisation. Initially, these antibodies were evaluated for their ability to neutralise an authentic SARS-CoV-2 strain. A strong correlation was observed between the pseudovirus and real virus neutralisation potencies (Fig. S2B). 16/19 ACE2 blocking antibodies (Fig. 3B, Fig. S3 and Table S4) neutralised the authentic virus with IC_50_ ranging from 100 pM (15 ng/ml) to ~670 pM (100 ng/ml). The control antibodies COV2-2196, COV2-2130 and S309 neutralised the virus with IC_50_s of 120 pM, 840 pM and 2.4 nM respectively. The two non-blocking neutralisers (ION_321 and ION_300) showed IC_50_s of 4.15 nM and 16 nM respectively. Finally, 4 potent antibodies were tested as Fabs and IgGs to determine the importance of valency in virus neutralisation. The neutralisation mediated by the IgGs were orders of magnitude more potent than that of equivalent Fabs (Fig. 3C), suggesting the likely bivalent engagement of both IgG arms. The increased virus neutralisation potency of IgG is likely a result of enhanced viral engagement due to avidity but could also include crosslinking spike proteins leading to aggregation of virions *(12)*.

Given the prevalence of RBD mutations among the circulating SARS-CoV-2 isolates, it is desirable to use a therapeutic approach that includes a cocktail of two or more antibodies directed against non-competing epitopes with different mechanisms of viral neutralisation. In addition to mitigating the risk of escape mutants, such an approach could also induce synergetic neutralisation effects *(13, 14)*. Therefore, the final panel of 21 neutralising antibodies were subjected to an epitope binning experiment using Octet Bio-Layer Interferometry to identify binding sites on RBD and their inter-relationships. Two control antibodies (COV2-2130 and COV2-2196) were also included in the analysis. Based on the pairing patterns, 7 different epitope bins were identified (Fig. 3D, Fig. S4). The majority of ACE2 blocking antibodies clustered into Bin A, which overlapped with 4 other bins. However, within this group, Bin B/Bin E combination and Bin C/Bin E combination were non-competitive. As expected, Bin G containing the two non-ACE2 blocking neutralisers, showed no overlap with any of the other bins, hence creating 5 additional pairings involving an antibody from each bin. Based on this, 10 combinations covering all non-competitive bins (except for antibodies from Bin D and F) were tested to identify antibodies that can be paired together in an antibody cocktail. Of the 10 combinations tested, only two combinations showed moderate synergy while 2 combinations were additive, and 6 combinations were antagonistic (Table 1). Importantly, ION_300 (a non-ACE2 blocker) from Bin G paired well with antibodies from other bins and was part of the two combinations that showed moderate synergy.

**Table 1.**
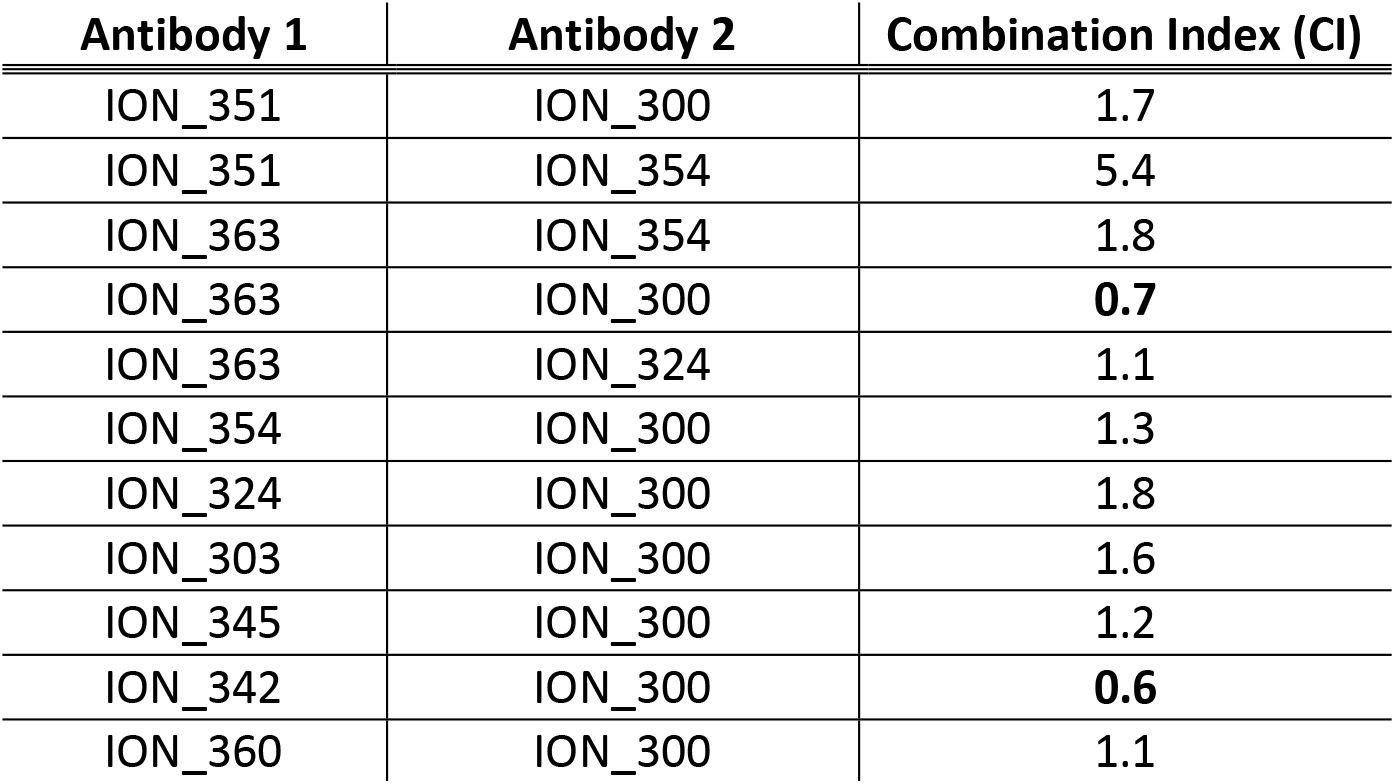
Effect of antibody combinations in the neutralisation of authentic SARS-CoV-2. Each antibody in the combinations listed was tested individually or at 1:1 ratio with the pairing antibody in the authentic SARS-CoV-2 neutralisation assay. The Combination Index (CI) for each antibody pair was calculated with CompuSyn *(39, 40)*. Based on the recommended cut-off values by CompuSyn, CI values <0.9 indicates synergy (**in bold**); 0.9<CI<1.1 indicate additive effect; CI>1.1 indicates antagonism. Antibody pair ION_351 and ION_354 from overlapping bins (A and C) was used as negative control for the assay.

| Antibody 1 | Antibody 2 | Combination Index (CI) |
| --- | --- | --- |
| ION_351 | ION_300 | 1.7 |
| ION_351 | ION_354 | 5.4 |
| ION_363 | ION_354 | 1.8 |
| ION_363 | ION_300 | **0.7** |
| ION_363 | ION_324 | 1.1 |
| ION_354 | ION_300 | 1.3 |
| ION_324 | ION_300 | 1.8 |
| ION_303 | ION_300 | 1.6 |
| ION_345 | ION_300 | 1.2 |
| ION_342 | ION_300 | **0.6** |
| ION_360 | ION_300 | 1.1 |

Biophysical characterisation of early-stage therapeutic candidates is important to identify any associated liabilities or risks that may hinder the progression of an antibody towards Chemistry, Manufacturing and Controls (CMC) *(15)*. Although the functional binding properties set the threshold for progressing antibodies into preclinical and clinal development, good thermal, physical and chemical properties (collectively known as “developability” properties) are all required to ensure they can be produced at scale with minimal loss and are tractable as a potential human drug. Sub-optimal developability properties can cause poor *in vivo* efficacy, PK/PD and immunogenicity leading to expensive late-stage clinical failures or high costs of manufacturing *(16, 17)*. Therefore, the final panel of neutralising antibodies were subjected to a series of experiments to determine the developability profile (see Supplementary Materials). This includes a pH stress test (to mimic the virus inactivation step during manufacturing), thermal stress test, freeze-thaw test, fragmentation analysis using CE-SDS, purity and column interaction test using HPLC-SEC, propensity for self-aggregation using AC-SINS and determination of isoelectric points using capillary isoelectric focusing (cIEF). 13 out of 21 (60%) antibodies passed the cut-off values for each assay and were hence deemed developable (Table S5). This represents an attrition rate of 40% from the initial panel of 21 neutralising antibodies tested, underlining the importance of developability testing as part of early antibody drug discovery.

### Crystal structures of ACE-2 blocking and non-blocking antibodies in complex with RBD

To understand and characterise the molecular basis of viral inhibition, we determined the crystal structure of the potent ACE2 blocking ION_360 bound to RBD and of the complementary non-receptor competitive ION_300 in complex with RBD. The structure of the ION_360-SARS-CoV-2 RBD complex was determined at 2.80 Å. Consistent with its antiviral potency, ION_360 binds to the ACE2 receptor binding motif (RBM) on the SARS-CoV-2 RBD, which is a feature reported for a number of antiviral antibodies *(10, 18)*. Examples of key interactions seen between the ION_360 antibody and the SARS-CoV-2 RBD are illustrated in Fig. 4A and Fig. S5.

**Fig. 4.**
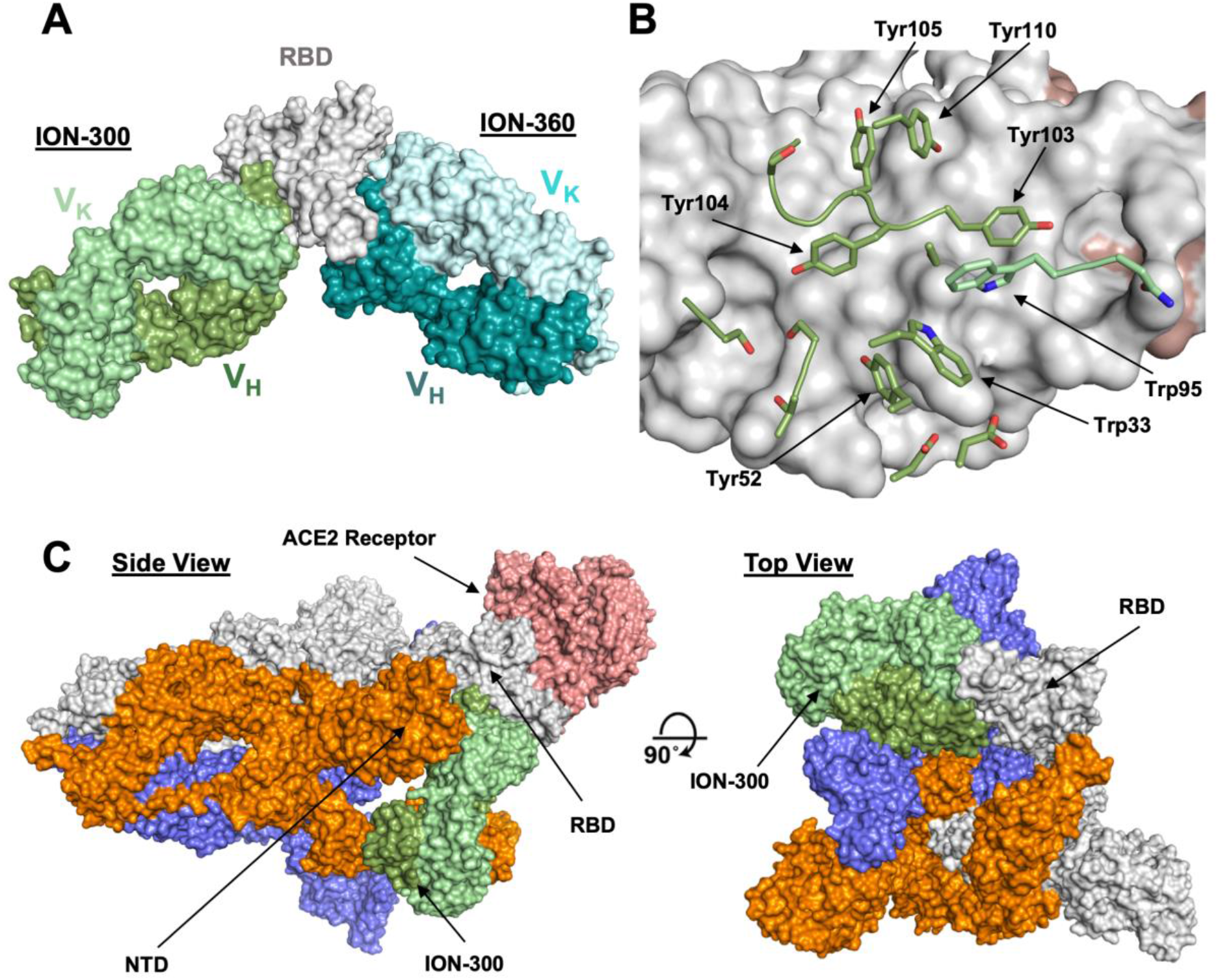
Structures of ION_300 and ION_360 antibodies bound to SARS-CoV-2 RBD. A) Molecular surface representation of ION_300 (greens) and ION_360 (blues) overlayed by their bound RBD (grey). B) Analysis of the ION_300:RBD interface. CDR residues within 5 Å of the RBD are shown in sticks for the V_H_ (green) and V_K_ (pale green) chains. The RBD is represented as a surface (grey) with ACE2 binding residues highlighted (salmon). C) Overlay of the ION_300:RBD complex (greens) onto a published cryo-EM structure of the spike protein trimer (grey, orange and blue) bound to the ACE2 receptor (salmon) (PDB: 7a96), shown in molecular surface representation. Figures prepared using PyMol.

To determine the mechanism of viral neutralisation of the non-ACE2 blocking antibody ION_300, the structure of the ION_300-SARS-CoV-2 RBD complex was solved by protein crystallography to 2.35 Å. Our structure revealed that the ION_300 interface with the RBD is predominantly through the V_H_ domain, burying 647 Å2 of antibody surface, compared to only 27 Å2 for the V_K_ domain. All three CDR loops from the V_H_ domain contribute at least three residues to the interface, whereas only two residues from the V_K_ domain are involved, with a total of 19 residues across both domains losing at least 10 Å^2^ of solvent accessibility following binding. The contact surface involves seven aromatic residues from ION_300 CDR loops, three of which are central to CDRH3 interactions (Tyr103, Tyr104, Tyr105), as well as seven hydrogen bonds and two salt bridges. An illustration of the RBD:ION_300 interface is shown in Fig. 4A and 4B. The structure of the complex reveals that ION_300 binds away from the receptor binding motif recognised by ACE2 on the RBD and against the exposed ß-pleated sheet, particularly impacting on residues K462-S469 immediately following the ß1’ strand.

It is important to note that the epitope identified for ION_300 is distinct from all current published anti-SARS-CoV-2 antibodies (Fig. S6). There is a very limited overlap with the epitope of Fab 52 (PDB: 7k9z *(19)*), which contacts the backside of the RBM, whereas ION_300 shows no contact with the RBM. Furthermore, the most prevalent recorded mutations, V367F, N439K, Y453F, S477N, V483A *(20)*, and the N501Y mutation associated with the recent virulent strain (B.1.1.7 lineage) identified in the UK, are not found within the epitope of ION_300 and are consequently considered highly unlikely to affect the antiviral potency of this antibody.

Structural alignment of the RBD from the ION_300 complex with the RBD from the ACE2 complex (PDB:6m0j) reveals very limited conformational differences, with an overall rmsd of 0.76 Å when aligning all Ca atoms, strongly suggesting that the antiviral mechanism of action of ION_300 is not allosteric. Furthermore, in reported cryo-EM structures of the SARS-CoV-2 spike protein, the ION_300 epitope on the RBD is buried behind the NTD of an adjacent S1 polypeptide chain when the RBD is in the closed conformation. The ION_300 binding site only becomes accessible when the RBD is in the open conformation (Fig. 4C), indicating that binding of this antibody may result in a spike protein which is locked in an RBD open type conformation. This perhaps suggests that ION_300 may utilise a similar ratcheting mechanism to that proposed by David Veesler’s Group *(21–23)*, in which the open/closed equilibrium of the spike protein, with regard to the RBD, is pushed to a fully open conformation following antibody binding. Such changes may cause a premature adoption of a post-fusion spike conformation, ultimately resulting in the inhibition of virus entry into host cells. Alternatively, steric hindrance or aggregation of virions via bivalent crosslinking of spike trimer could also contribute to viral neutralisation function of ION_300 *(12)*.

### RBD-binding antibodies arise from recent B cell activation and correlate with patient status

The BCR repertoire sequence data from these patients has been previously published *(24)*, yielding 3,485,995 unique VH sequences across all patients. To investigate the B cell responses of the patients in more detail, the sequence data from the 254 unique RBD-binding antibodies isolated by phage display were integrated with the total BCR repertoire data (Fig. 1). Co-clustering of the two datasets was performed using a previously described threshold (see Methods) to group together sequences that are sufficiently similar to be considered part of the same B cell clonal expansion, and likely targeting the same epitope. This clustering yielded 838,887 clusters across all of the patients; these clusters were then labelled according to whether they contained any of the RBD-binding antibodies identified from the phage-display process. Of the 254 unique RBD-binding antibodies, 201 of these co-clustered with the total BCR sequence data. Several of these 201 antibodies mapped to the same cluster, resulting in 89 different clusters that could then be annotated as RBD-binding. 108 of the 201 RBD binding antibodies also showed >30% neutralising activity in the pseudoviral neutralisation assay (at 25 nM), enabling 49/89 of the RBD-binding clusters to also be annotated as containing neutralising antibodies.

The mean size of all clusters across patients was 4 sequences, but the clusters annotated as containing RBD-binding or neutralising antibodies contained on average 116 and 63 sequences respectively, indicating that these B cells are undergoing clonal expansion (Fig. 5A). Investigating the isotype subclass distributions of the sequences within these clusters showed different distributions for total clusters, clusters annotated as RBD-binding, and clusters annotated as RBD binding and neutralising (a subset of RBD binders that neutralise the virus, Fig. 5B). 71% of RBD-binding clusters, and 80% of neutralising clusters contained IgM sequences, indicating recent activation of these B cells. Furthermore, calculating the mean mutation from germline of the sequences within the clusters showed that the RBD-binding and neutralising clusters had fewer mutations than total clusters (RBD-binding: 2.6, Neutralising: 2.2, Total: 7.6), giving further evidence that the RBD-binding and neutralising clusters have arisen from recently activated B cells (Fig. 5C).

**Fig. 5.**
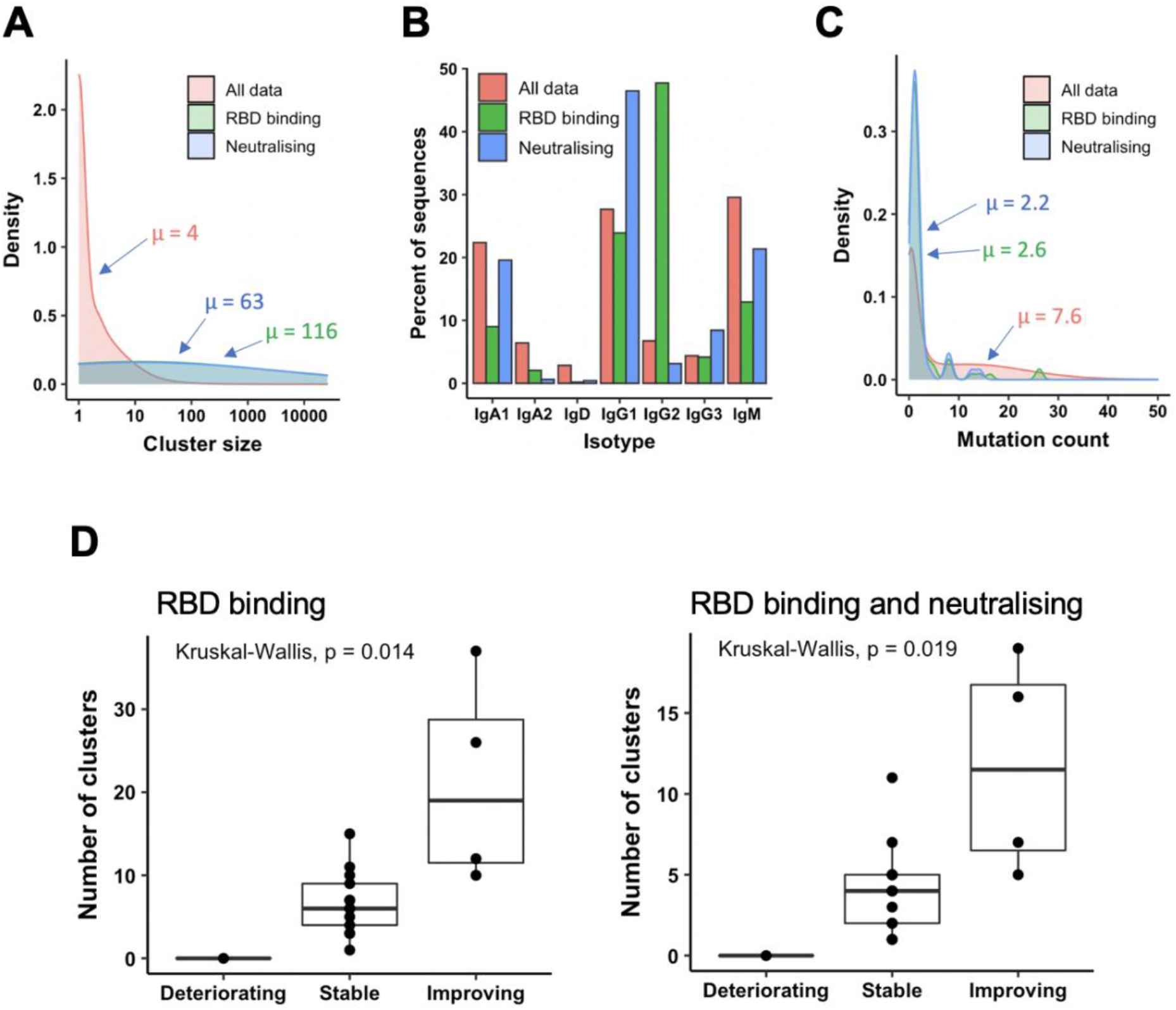
Relating antibodies discovered by phage display back to patient B cell repertoire data. A) Sequences clustered into related groups. Clusters then annotated based on whether they contained RBD-binding or its subset of neutralising antibody sequences. Density plot shows distribution of clusters of different sizes in the combined dataset from all 18 patients. B) The isotype subclass distribution of sequences belonging to the different groups of clusters. C) Mean mutation of all sequences within each cluster was calculated. Density plot shows the distribution of clusters with different numbers of mutations. D) The number of clusters annotated as RBD binding (left), or RBD binding and neutralising (right) in each patient, stratified according to disease status.

Looking at total clusters, there is a bimodal distribution of mutation counts, which becomes more pronounced when considering the clusters that do not contain IgM sequences (Fig. S7A), and those with evidence for clonal expansion (Fig. S7B). We interpret the first mode of the distribution to represent naïve or recently activated B cells, and the second mode to represent memory B cells. By cutting this bimodal distribution at 5, we classified whether the RBD-binding clusters had arisen from a naïve response (<5 mutations), or memory recall (>= 5 mutations). In total, 88% of the RBD-binding clusters, and 90% of the neutralising clusters were classified as having arisen from recent activation. We can thus conclude that the response to COVID-19 is largely driven by naïve B cell activation, and there is very little re-activation of circulating memory cells.

Next, the number of clusters annotated as either RBD-binding or neutralising was calculated independently for each patient. There were no RBD-binding clusters identified in the one deteriorating patient (Fig. 5D), but at least one identified in every stable and improving patient. There were significant differences in the number of both RBD-binding and neutralising clusters identified between each of the three patient groups. There were more RBD-binding and neutralising clusters identified in the improving compared to the stable, and the stable compared to the deteriorating groups. Improving patients therefore provide the best source for therapeutic antibody discovery; the deteriorating patients may have yet to mount an effective immune response.

### The antibody response to COVID-19 is highly convergent

We and others have previously observed a stereotypic BCR response to COVID-19 infection *(24)*. We replicate the findings here, showing that the clusters annotated as RBD-binding use a restricted set of V gene segments (Fig. S8). Clusters utilising VH3-30, VH3-53, VH3-66, VH3-9 and VH5-51 dominated the response, with lower levels of VH1-18, VH1-2, VH1-49, VH1-69, VH3-11, VH3-13, VH3-15, VH3-20, VH3-23, VH3-48, VH3-64 and VH3-7 also seen. Heavy and light chain V gene utilisation among the RBD binders identified from phage display screening is shown in Fig. S9.

We have now extended these observations by investigating the specific convergence between individuals of the clusters annotated as RBD-binding or neutralising. Of the 89 clusters annotated as RBD-binding, 26% (23/89) of the RDB binders and 33% (16/49) of the neutralising clusters were convergent across at least 2 individuals. This is in stark contrast to the convergence seen across the entire dataset which is 2.5% of total clusters. Furthermore, one of the RBD-binding and neutralising clusters was shared between 14 of the 18 different individuals in the study (Cluster ID 3, Table 2.1). Interestingly, the moderately neutralising non-ACE2 blocking antibodies were also abundant and convergent with one clonally expanded cluster (size = 128) recurring among 7 individuals.

**Table 2.**
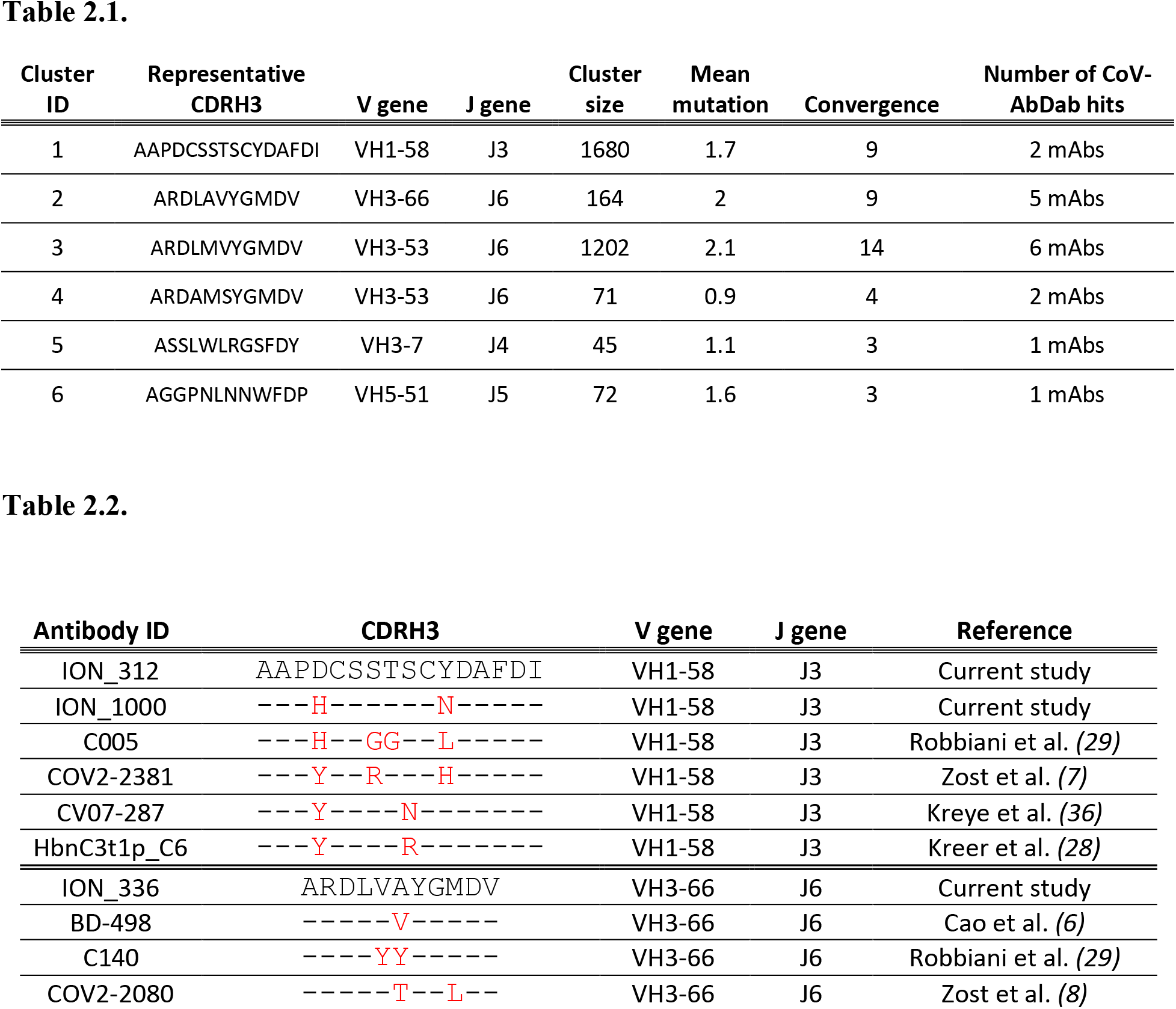
Convergent clusters identified in the current study which also had sequences map to them from the CoV-AbDab **(Table 2.1)**. This table shows the properties of convergent antibody clusters identified from the current study, including the size of the cluster (i.e. unique antibodies found in each cluster, an indicator of clonal expansion), number of different patients each cluster was present in (convergence) and number of antibodies found in the CoV-AbDab database with the same cluster identity. Convergence of antibody sequences across separate studies **(Table 2.2)**. Heavy CDR3 sequence similarity (>80%), V gene, J gene usage shown between antibodies identified within the current study to those in separate published studies.

In addition to investigating the convergence between individuals in the current study, we also investigated the convergence with SARS-CoV-1/2 binding antibodies identified in other studies. We extracted all V_H_ sequences from the CoV-AbDab *(25)* annotated as having a human-derived V gene segment, and having V and J gene segment annotations, and CDRH3 sequence determined. This yielded 1,051 unique sequences. These sequences were then also integrated into the clustering analysis of phage-derived RBD-binding and neutralising antibodies, and the total BCR repertoire data. In total, 112 of the clusters in the total BCR repertoire contained antibodies that had been extracted from the CoV-AbDab, showing that convergence is seen between studies. Furthermore, there were six clusters which contained both antibodies from the CoV-AbDab and antibodies from the current study (Table 2.1). All of these clusters were classified as having arisen from recent activation based on having <5 mean mutations, and the presence of IgM sequences. These clusters were all convergent between at least three individuals in the current study. Two of these clusters (Cluster ID1 and Cluster ID2) were also convergent between nine individuals in this study, as well as three separate studies from the CoV-AbDab. Both of these clusters were in the panel of the 21 most potent antibodies discovered in the current study. For example, antibodies ION_312 and ION_1000, were highly similar to a clonally expanded sequence cluster (containing 1680 unique sequences, Cluster ID1, Table 2.1) from 9 different patients in this study and antibodies from 4 separate studies (Table 2.2), indicating its importance in mediating protection.

## Discussion

We have previously described *(24)* B cell receptor repertoire analysis of the V_H_ populations from 18 SARS-CoV-2 positive donors at an acute stage of disease. In the present study we have constructed libraries from the antibody repertoire of these patients and used phage display to identify a sub-set of antibody genes which encode functional binders and potent neutralisers of SARS-CoV-2. Unlike most direct B cell screening methods, phage display technology allows processing of millions of antibody genes from large numbers of patient donors. It also allows enrichment of functional clones that may be rare or have not yet undergone clonal expansion within the initial antibody response. This deep mining approach yielded neutralisers to diverse epitopes within the RBD. This includes a group of highly potent antibodies which compete with ACE2 for RBD binding, with neutralisation IC_50s_ matching the best antibodies reported with similar mechanism of action *(5, 7, 8, 26, 27)*.

Another group included antibodies with moderate potencies which neutralised the virus through a distinct mechanism of action (independent of inhibiting the RBD-ACE2 interaction) including an antibody to a unique epitope (ION_300). Despite their relatively weaker potency, non-ACE2 blocking neutralisers were abundant within the clonally expanded patient antibody response with strong convergence scores, indicating that such antibodies also play a key role in providing protection. The crystal structure of ION_300 revealed that it binds to a unique epitope at the opposite side of the RBM. Identification of contact residues within the epitope suggest that it is likely to retain binding to widely circulating RBD mutations including the N501Y mutation that is predicted to enhance the infectivity of the newly identified SARS-CoV-2 B.1.1.7 lineage. In addition, this antibody paired well with potent antibodies from ACE2-competing epitope bins to impart synergy (albeit moderate), providing a compelling case for inclusion in an antibody cocktail for in vivo evaluation or further potency maturation.

Sequence analysis of the selected functional binders from this study shows that many of the potent antibodies are identical to, or within a few amino acids of, germline encoded V and J sequences. This is in line with previous reports from Kreer et al, who describe isolation of neutralising IgG antibodies with a spectrum of variable domains with low levels of somatic mutation derived from memory B cells *(28)*. Projection of the sequences of the confirmed RBD binders and neutralisers identified in this study onto a dataset derived from the same patient group including VHs from both IgG and IgM populations, confirms that a number of these potent neutralisers (originally selected from the IgG pool) are in fact represented in the IgM repertoire. Thus, the occurrence at an early stage of infection of relatively unmutated V_H_ genes within both IgM and IgG populations is indicative of recently class-switched B cells that have yet to go undergo somatic hypermutation.

By combining the identification of highly potent neutralising antibodies with deep sequencing we highlight convergence within the antibody response among different patients within this study group and beyond. The occurrence of a convergent antibody response among COVID-19 patients is described to a lesser extent by other studies. Robbiani et. al. purified IgG-expressing, RBD-binding memory B cells from a limited number of convalescent COVID-19 patients (n=6) at an average of 39 days post-symptom onset *(29)*. Analysis of the antibody genes revealed the presence of closely related antibodies in different individuals at this later time point in the memory pool. Here, we show evidence of recurring antibody genes within the total antibody pool (including the IgM repertoire) of 18 patients and also among the published anti-SARS-CoV-2 antibody sequences from multiple independent studies, including one of the most convergent sequences from Robbiani et al (Table 2.2).

Convergent antibody responses have been described in the response to other infectious diseases *(30–32)*, but to our knowledge, the extreme level of convergence in the COVID-19 response has not been reported in other disease settings. The occurrence of potent neutralising antibodies within the germline encoded naïve repertoire may be part of the explanation. The high convergence of specific BCR sequences in COVID-19 that have protective properties suggests that developing these into antibody therapeutics could be highly effective. Monitoring for the development of these sequences may also be used as a generic method for assessing efficacy of novel vaccine strategies.

The majority of characterised antibodies from these patient derived libraries had affinities in the range of 1-30 nM, which is typical of antibodies isolated from IgM-derived naïve phage display repertoires *(33–35)* and did not show any strong correlation with neutralisation potency. The ability of a naïve-like antibody response with moderate binding affinities to impart highly potent viral neutralisation could be rationalised by the avid binding of bivalent IgGs to the trimeric spike protein (the main target of neutralising antibodies reported here or elsewhere). The importance of valency in neutralisation is demonstrated by the comparison of neutralisation IC_50_s of IgGs and Fabs (Fig. 3C).

While the devastating effect of SARS-CoV-2 infection on the population is clear to see, this study clearly demonstrates the presence of potent neutralising antibodies within the armoury of the naïve repertoire protecting the majority of the population. This readiness may be explained by the combination of high diversity derived from the antibody germline locus together with a multivalent presentation of antibodies effecting potent neutralisation of polyvalent targets.

## Materials and Methods

### Recombinant antigen and control antibodies

For expression of the RBD subdomain of the SARS-CoV-1 (residues Arg306 to Phe527) and SARS-CoV-2 (residues Arg319 to Phe541) spike protein and human ACE2, genes encoding the proteins were synthesised and cloned upstream of either an Fc tag or rCD4-Avi tag, both with an additional 6 × His tag in mammalian expression vectors *(37)*. The constructs were expressed in Expi293F™ cells (Thermo Fisher, A14527) and purified by affinity chromatography using Nickel-NTA agarose (Invitrogen, R90115). SARS-CoV-2 S1 protein with a His tag (S1N-C52H3) and SARS-CoV-2 S protein active trimer with His tag (SPN-C52H8) were separately purchased from ACRO biosystems. MERS-CoV S1 with a His tag was purchased from Sino Biological (40069-V08H).

All antigens with an Avi tag were biotinylated enzymatically using BirA biotin-protein ligase (Avidity, Bulk BirA) while non-Avi tagged antigens were biotinylated chemically using EZ-Link Sulfo-NHS-Biotin (Thermo Fisher, A39256). V_H_ and V_L_ sequences of SARS-CoV-2 control antibodies (COV2-2196, COV2-2130, S309, B38, H4, and CR3022) were obtained from the CoV-AbDab database *(25)* and cloned into pINT3/pINT54 IgG expression vectors *(4)* as synthetic gene fragments. The antibodies were expressed using the Expi293™ system and purified using Protein-A affinity chromatography (Generon, PC-A100). Control antibody SAD-S35 was purchased from ACRO biosystems.

### Sample collection and total RNA preparation

Peripheral blood was obtained from a subset of 18 patients who were admitted to medical wards at Barts Health NHS Trust, London, UK with acute COVID-19 pneumonia, after informed consent by the direct care team. Blood collection and processing, followed by total RNA preparation, was carried out as previously described *(24)*. RNA was split for use in both construction of phage libraries (see Supplementary Materials) and BCR sequencing.

### Phage display panning, subcloning and high throughput IgG expression

The hybrid and fully patient derived libraries (see Supplementary Materials) were subjected to three rounds of phage display selections against biotinylated RBD-Fc-His, RBD-rCD4-Avi-His, and S1-His were carried out as previously described *(2)*. The first and second rounds used selection antigen concentrations of 50 nM and 5 nM respectively. In the third round, each selection was performed at two selection antigen concentrations, 0.25 nM and 0.05 nM. Antibody genes were isolated from the third-round selection populations and sub-cloned into the pINT3 and pINT54 IgG1 mammalian expression vectors *(4).* Transfection-quality plasmid DNA was prepared for 1498 clones in 96 well plate format (Biobasic, BS415). The DNA was then expressed by transient transfection in Expi293F™ cells (Thermo Fisher, A14527) in 96 well plates at 700 μl/well scale.

### Primary screening and sequencing

An affinity capture assay was performed as the initial screening of the 1,498 expressed IgGs. The assay was performed as described elsewhere *(3)* using 1 nM biotinylated RBD-rCD4-His or S1-His. V_H_s and V_L_s were sequenced using Eurofins Genomics and Genewiz Sanger sequencing services. To identify unique combinations of heavy CDR3 and light CDR3, antibody frameworks and CDR regions were annotated and analysed using Geneious Biologics (Biomatters).

### SARS-CoV-2 RBD-ACE2 blocking assay

Black 96-well immune plates (Thermo Scientific, 10030581) were coated with mouse anti-rCD4 antibody (Bio-Rad, MCA1022R) at 4 °C overnight. The following day, plates were blocked with 3% (w/v) milk powder (Marvel) in PBS, before incubation with purified recombinant SARS-CoV-2 RBD-rCD4-His (1.25 μg/mL) for 1 hour. To identify antibodies that could block the interaction between RBD and ACE2, the anti-SARS-CoV-2 antibodies were pre-incubated with 2 nM of recombinant ACE2-Fc-His-Biotin for 30 minutes before transferring to the RBD-coated plates. The IgGs and ACE2 were incubated in the plates for 1 hour and ACE2-Fc-Biotin was detected by DELFIA Eu-N1 Streptavidin (Perkin Elmer, 1244-360) using the DELFIA-TRF system (Perkin Elmer).

### SARS-CoV-1 and MERS-CoV cross-reactivity assay

Black 96-well immune plates (Thermo Scientific, 10030581) were coated with SARS-CoV-1 RBD-rCD4-His or SARS-CoV-2 RBD-rCD4-His or MERS-CoV S1-His (Sino Biological, 40069-V08H), at 3 μg/mL, at 4 °C overnight. The following day, plates were blocked with 3 % (w/v) milk powder (Marvel) in PBS before incubation with purified IgGs (2 μg/mL) for 1 hour. Bound IgG was detected with Eu-N1 Anti-Human IgG (Perkin Elmer, 1244-330). using DELFIA-TRF system (Perkin Elmer).

### Neutralisation assays and combination testing

Briefly, a lentiviral pseudotyped virus was used for the infection of HEK293T/17 cells transiently expressing human ACE2 and TMPRSS2 for the pseudoviral assay. The SARS-CoV-2 Australia/VIC01/2020 isolate (Centre For AIDS Reagents cat. no. 100980) was used to infect VERO CCL-81 cells for the authentic virus neutralisation assay. The Combination Indexes (CI) for the antibody combinations were calculated using CompuSyn based on Chou-Talaley method *(39, 40)*. See Supplementary Materials for detailed methods.

### Epitope binning and affinity measurement

The epitope binning of anti-SARS-CoV-2 IgGs was carried out using Octect BioLayer interferometry using the classical sandwich format (Fig. S4). Surface Plasmon Resonance (SPR) experiments were performed using the Sierra SPR-32 instrument (Bruker). 1:1 binding kinetics (at 25°C) were determined by flowing different concentration of monomeric SARS-CoV-2 RBD over antibodies immobilised on biosensor chips. See Supplementary Materials for detailed methods.

### Expression and purification

For medium scale expression for detailed characterisation, all antibodies were expressed at 50 ml scale in ExpiCHO or Expi293 system (Thermo Fisher). Expressed antibodies were purified using protein-A affinity chromatography (by HiTrap Fibro PrismA unit, Cytiva, 17549855) followed by size exclusion chromatography (HiLoad Superdex 200 16/600, Cytiva, 28-9893-35).

For single concentration pseudovirus assay, SPR, and cross-reactivity evaluation, the antibodies expressed in 96 well plates (Expi293 cells) were purified by protein-A affinity chromatography (Generon, PC-A100), using 96 well filter plates (Whatman Polystyrene Unifilter Microplates, GE Healthcare, 11313535). After purification, antibodies were buffer exchanged into PBS using Zeba™ Spin Desalting Plates, 7K MWCO (ThermoFisher Scientific, 89807).

### B cell receptor repertoire sequencing

Three different V_H_ sequence datasets were integrated for analysis: 1) The V_H_ sequence data from the antibodies isolated in the present study using phage display, 2) the B cell receptor repertoire V_H_ sequence data generated from the same patients and previously published *(24)* and 3) all V_H_ sequences in CoV-AbDab [accessed 28^th^ August 2020] annotated as having a human-derived V gene segment, and having V and J gene segment annotations, and CDRH3 sequence determined.

Each sequence from the combined dataset was processed using IgBlast *(42)* to determine V and J germline gene segment usage, and locations of the CDRs and FWRs. Mutation count was determined by the number of mismatches between the sequence and its inferred germline using the shazam R package *(43)*. For dataset 2, the isotype of each sequence was possible to determine by comparison to germline constant region sequences. For datasets 1 and 3, this information was not available.

Sequences were clustered into groups using a previously described algorithm *(24)*. This is a greedy clustering algorithm, run with a threshold requiring sequences within a cluster to have the same V and J gene segment, and no more than 1 AA mismatch per 10 AAs in the CDRH3. This threshold has been previously shown to group together sequences that are sufficiently similar to be considered part of the same B cell clonal expansion, and likely targeting the same epitope. The cluster centre is defined as the most common sequence within the cluster. All datasets, and all samples within each dataset were clustered together to identify overlap between the datasets.

### Protein expression and purification for crystallography studies

The Fab fragments for anti-SARS-CoV-2 antibodies ION_300 and ION_360 and SARS-CoV-2 receptor binding domain (RBD) were expressed in Expi293F™ cells (Thermo Fisher, A14527). The antibody:RBD complexes were mixed and co-purified by size exclusion chromatography in 20 mM Tris-HCl (pH 7.5) and 50 mM NaCl. For crystallisation, antibody:RBD complex samples were concentrated to 5-10 mg/mL.

### Crystallization, structure determination, and refinement

All crystals were obtained by the vapor diffusion method at 19°C, by mixing equal volumes of protein plus well solution. The RBD:ION_300 crystals grew in 16% PEG3350 and 0.3 M potassium citrate tribasic, whereas crystals of RBD:ION_360 crystals grew in 16% PEG3350 and 0.2 M ammonium citrate tribasic. For cryoprotection, crystals were generally transferred to a solution of mother liquor plus 22% ethylene glycol. Data sets were collected at the European Synchrotron Radiation Facility, Grenoble (beamline ID30A1 (MASSIF-1)). Data from crystals of RBD:ION_300 and RBD:ION_360 were refined to 2.35 and 2.80 Å resolution, respectively. Data were processed using XDS *(44)* and AIMLESS from the CCP4 Suite *(45)*. Cell parameters and data statistics are summarized in Table S6.

Both complex crystal structures were solved by molecular replacement using Phaser *(46)* utilising the coordinates from the SARS-CoV-2 RBD domain (PDB: 7JMP) and using homology models for ION_300 and ION_360 antibodies generated using SWISSMODEL *(47)* to model the heavy and light chains. Atomic models were built using Coot *(48)* and refined with Refmac *(49)*. All structures were solved by molecular replacement and are reported with final Rwork/Rfree values below 20/25% with good stereochemistry (Table S6.).

## Supplementary Materials

Detailed materials and methods for library construction, neutralisation assays and biophysical characterisation of anti-SARS-CoV-2 antibodies.

**Fig. S1.** Dose-response curves demonstrating pseudovirus neutralisation.

**Fig. S2.** Correlation of pseudovirus neutralisation with authentic virus neutralisation and affinity.

**Fig. S3.** Dose-response curves demonstrating authentic virus neutralisation.

**Fig. S4.** Epitope binning with RBD on panel of 21 antibodies and controls.

**Fig. S5.** Analysis of the ION_360:RBD Interface.

**Fig. S6.** Superposition of antibody and nanobody structures targeting the SARS-CoV-2 RBD.

**Fig. S7.** Mutation distribution of clusters.

**Fig. S8.** VH germline usage in convergent antibody response.

**Fig. S9.** Heavy and light chain V gene utilisation and pairing preference of all RBD-binding antibodies.

**Table S1.** Days of symptoms prior to sample collection across related studies.

**Table S2.** Clinical details and characteristics of the patients diagnosed with SARS-CoV-2 whom blood was collected from for use in this study.

**Table S3.** 1:1: binding kinetics of the final panel of 21 antibodies to SARS-CoV-2 RBD measured by SPR.

**Table S4.** Neutralisation potency of the selected 21 antibodies against a pseudotyped virus and an authentic strain of SARS-CoV-2.

**Table S5.** Summary of developability data for final panel of 21 antibodies.

**Table S6.** X-ray data and refinement statistics.

## Acknowledgements

The authors wish to thank the UK Bioindustry Association (BIA) antibody taskforce, in particular, Deirdre Flaherty for making the collaboration between the contributing organisations possible.

## Author contributions

A.K-V, J.Mc, J.O, J.G conceived the study. P.P and W-Y.L recruited the patients and executed clinical protocols. A.K-V, K.S, J.G, L.M, P.V, S.S, M.C, S.H, D.M, J.C, G.M and O.A designed experiments. G.B, P.V, L.M, L.L, E.M, K.P, K.C, D.P, L.C, C.T, I.D carried out the antibody selection and screening. R.L, P.V, E.M, D.T, S.M, R.B carried out expression, purification and biophysical characterization. G.M and E.B performed neutralization assays. G.H and M.B performed the structural studies. A.K-V, G.B, J.G, J.Mc, L.L, L.M, G.H and K.S wrote the manuscript.

## Data and materials availability

Materials and sequence generated in this study will be made available on request, but we will require a completed Materials

Transfer Agreement signed with IONTAS Ltd.

