## Supplementary figures and tables for "Deep mining of early antibody response in COVID-19 patients yields potent neutralisers and reveals high level of convergence"

**This PDF file includes:**

Detailed materials and methods for library construction, neutralisation assays and biophysical characterisation of anti-SARS-CoV-2 antibodies.

**Fig. S1.** Dose-response curves demonstrating pseudovirus neutralisation.

**Fig. S2.** Correlation of pseudovirus neutralisation with authentic virus neutralisation and affinity.

**Fig. S3.** Dose-response curves demonstrating authentic virus neutralisation.

**Fig. S4.** Epitope binning with RBD on panel of 21 antibodies and controls.

**Fig. S5.** Analysis of the ION_360:RBD Interface.

**Fig. S6.** Superposition of antibody and nanobody structures targeting the SARS-CoV-2 RBD.

**Fig. S7.** Mutation distribution of clusters.

**Fig. S8.** VH germline usage in convergent antibody response.

**Fig. S9.** Heavy and light chain V gene utilisation and pairing preference of all RBD-binding antibodies isolated using phage display technology.

**Table S1.** Days of symptoms prior to sample collection across related studies.

**Table S2.** Clinical details and characteristics of the patients diagnosed with SARS-CoV-2 whom blood was collected from for use in this study.

**Table S3.** 1:1: binding kinetics of the final panel of 21 antibodies to SARS-CoV-2 RBD measured by SPR.

**Table S4.** Neutralisation potency of the selected 21 antibodies against a pseudotyped virus and an authentic strain of SARS-CoV-2.

**Table S5.** Summary of developability data for final panel of 21 antibodies.

**Table S6.** X-ray data and refinement statistics.

### Materials and Methods

#### Library construction

Two library strategies were employed, both using the antibodies in an scFv format with variable heavy chain ($V_H$) and variable light chain ($V_L$) fragments joined by a flexible linker peptide ($Gly_4$ Ser $Gly_4$ Ser $Gly_3$ Ala Ser). The first strategy was to construct hybrid libraries where donor $V_H$s were combined with a library of naïve kappa and lambda $V_L$s, whereas the second strategy was to construct a complete patient derived library by assembling $V_H$s and $V_L$s on a donor-by-donor basis.

Antibody $V_L$ and $V_H$ repertoires were obtained through cDNA synthesis from total RNA using First-Strand cDNA synthesis kit (GE Healthcare, 27926101), followed by PCR amplification. The cDNA template for $V_H$s were synthesised using an IgG-specific primer (5'-AGTAGTCCTTGACCAGGCAG -3') and the cDNA template $V_L$s were synthesised using a pd$(N)_6$ random hexamer primer to allow for amplification of both lambda and kappa sequences. For the hybrid library strategy, the $V_H$s from the cDNA template were amplified and cloned into naïve kappa and lambda light chain libraries as described before *(2)*.

For the complete patient derived library approach, the $V_H$s and $V_L$s (both lambda and kappa) from each donor were amplified and assembled on a donor-by-donor basis. The primary amplification of $V_H$s for the hybrid libraries was also used as the first step for the generation of $V_H$s of the complete patient derived libraries. Assembled fragments were cloned into the phage display vector pIONTAS1 using NcoI/NotI cloning sites *(2)*. Final libraries were obtained by electroporation of electrocompetent TG1 cells (Lucigen) with the purified ligation products.

### Pseudovirus neutralisation assay

To generate SARS-CoV-2 lentiviral pseudotyped virus, 5 x $10^6$ HEK293T/17 cells were seeded within a 10 cm dish. The following day, plasmids encoding the HIV-1 *gag-pol* genes (p8.91), a firefly luciferase reporter gene (pCSFLW) and the SARS-CoV-2 Spike gene (pCAGGS SARS-2-Spike) were transfected concurrently at a ratio of 1:1.5:1 µg, respectively, using Fugene-HD (Promega) transfection reagent. After overnight incubation, culture media was replenished. Supernatant was harvested at 48- and 72-hours post-transfection and stored at -80$^o$C.

For the assay, target cells were prepared by transient transfection of HEK293T/17 cells with 2 µg ACE-2 and 150 ng TMPRSS2 expression plasmids using Fugene-HD (Promega) transfection reagent. 24 hours post-transfection cells were detached and seeded at 20,000 cells/well within a 96-well plate and incubated at 37 $^o$C and 5% $CO_2$ for at least 2 hours. Three-fold serial dilutions of antibodies were prepared within 60 µL DMEM supplemented with 10% FBS and 1% penicillin/streptomycin. To each well, 60 µL containing 200 TCID50 (50% Tissue Culture Infective Dose) of SARS-CoV-2 lentiviral pseudotyped virus was added and incubated for 1 hour at 37 $^o$C. Following incubation, 100 µL of the purified antibodies and pseudotyped virus mix was transferred to the target cells and incubated for 72 hours at 37 $^o$C and 5% $CO_2$. To acquire results, luciferase expression was detected using the Promega Bright-Glo™ assay system and GloMax Navigator plate reader, following manufacturer's instructions. The half maximal inhibitory concentration ($IC_{50}$) was calculated using GraphPad version 8 as previously described *(38)*.

### Authentic SARS-CoV-2 virus neutralisation assay

VERO CCL-81 cells were seeded at 20,000 cells/well in a 96-well plate in DMEM (Sigma, D6546) supplemented with 2 mM L-Glutamine, 10 % Foetal bovine serum (PAN

Biotech UK) and 1% penicillin or streptomycin. The following day, two-fold serial dilutions of the antibodies in 60 µL of serum free DMEM were added to the same volume of media containing 100 $TCID_{50}$ of SARS-CoV-2 Australia/VIC01/2020 isolate (Centre For AIDS Reagents cat. no. 100980) and incubated for 1 hour at 37°C. The media from the VERO CCL-81 cells was removed and 100 µL of the antibody and virus mix was added to each well. 100 µL of DMEM supplemented with 4% FBS was added to each well. After 48 hours, cells were fixed with PBS containing 4% formaldehyde in PBS, for 1 hour at room temperature. Cells were permeabilised with 0.1% Triton X-100 in PBS for 15 minutes at room temperature. Cells were blocked with 3% milk in PBS containing 0.05% tween-20. Virus infection was detected using an HRP-conjugated anti-N protein antibody (The Native Antigen Company, MAB12184-HRP) for 1 hour at room temperature. Signal was detected using TMB substrate and stopped with $H_2SO_4$. The half maximal inhibitory concentration ($IC_{50}$) was calculated using GraphPad version 8 as previously described *(38)*.

Effect of the antibodies combinations was evaluated in the neutralisation assay by combining two antibodies at the ratio 1:1. Two-fold serial dilutions of the antibodies individually or in combinations were added to 100 $TCID_{50}$ of SARS-CoV-2, Australia/VIC01/2020 isolate. Assay was performed as described above. Response to each antibody concentration was normalised to the virus only value (0% neutralisation) and cell only value (100% neutralisation). The Combination Index (CI) for the antibodies combinations was calculated using CompuSyn *(39, 40)*.

**Epitope binning**

The panel of anti-RBD human antibodies was segregated into epitope bins using the classical sandwich Octet format (Fig. S4) on Anti-Human Fc Capture (AHC) biosensors (Sartorius, 18-5060). The experimental workflow was carried out in cycles as follows: Load of

Ab1 (5 minutes; 5 µg/ml), Quench/Block with irrelevant human IgG (5 minutes; 50 µg/ml; Jackson ImmunoResearch, 009-000-003), Wash in PBS (2 minutes), Load of SARS CoV-2 RBD (5 minutes; 5 µg/ml; Acro BioSystemsm, SPDC52H3), Baseline in PBS (2 minutes), Association of Ab2 (5 µg/ml; 5 minutes), Dissociation in PBS (2 minutes). Before and after each cycle, PBS-hydrated AHC biosensors underwent of 3x5-second cycles of regeneration (10mM Glycine pH 1; Sigma; G7403) and neutralisation (PBS). Reagents (200 µl/well) dispensed into black non-binding 96-well plates (Greiner; 655209) were maintained at 30°C for 5 minutes prior to and throughout the duration of the experimental cycles. Each antibody was investigated in both orientations (i.e. Ab1 vs Ab2). Irrelevant human IgG as Ab2 served as a control for reference subtraction. The data was acquired on an Octet HTX instrument using the 16 channels-high sensitivity mode with shaking (1000rpm). Software programs used were Octet Acquisition software (version 11.1) and Octet Data Analysis HT (version 11.1).

**Affinity determination using Surface Plasmon Resonance**

Surface Plasmon Resonance (SPR) experiments were performed using the Sierra SPR-32 instrument (Bruker). All measurements were performed at 25 ºC. A high-capacity amine sensor (Bruker, 1862614) was activated using EDC/NHS, before protein G was added at 150 µg/mL with a flow rate of 15 µL/min for 400 seconds resulting in immobilization of 3,000-5,000 RU per sensor spot. The sensor spots were inactivated using ethanolamine. IgG antibodies at 5 nM concentrations were captured at 15 µL/min for 180 seconds resulting in an average response of 100-200 RU. SARS-COV2 RBD-rCD4 at concentrations ranging from 20 nM to 0.625 nM was injected at 30 µL/min for 120 seconds and dissociation was recorded for 600 seconds. Following dissociation, the sensors were regenerated using 10 mM glycine-HCl, pH 1.5. The initial kinetics were conducted with 2 analyte concentrations used for each antibody. Measurements for the final panel of 21 antibodies were conducted in triplicates and for determination of kinetic constants, at least 4 analyte concentrations were used for each antibody.

### Developability assessment of anti-SARS-CoV2 antibodies

#### pH stress test

Affinity-captured samples were eluted from a 0.4 ml HiTrap Fibro PrismA unit using 0.1 M sodium citrate (pH 3.5), held for at least 30 minutes (virus inactivation) and neutralised using 1 M Tris-HCl (pH 9) in a 4 : 1 (sample: Tris-HCl) ratio. Analytical samples (10 µl) were taken and injected onto a Superdex 200 Increase 5/150 (Cytiva) using an Agilent 1260 Infinity HPLC system. Detection of aggregates was performed using a multi-angle light scattering (MALS) detector (DAWN HELEOS; Wyatt technology) and refractometer (Optilab TRex; Wyatt technology) coupled in-series to the UV detector. Peak analysis was performed using the software, Astra (Wyatt technology).

#### Determination of melting temperature

Melting temperatures (Tm) were determined by monitoring changes in intrinsic fluorescence (300 – 430 nm) versus temperature using the Uncle (Unchained labs). 9 µl samples (1 mg/ml) in PBS were thermally ramped from 25°C to 95°C (0.5 °C/min) and measured in triplicate. Analysis of the fluorescence signal was performed using the barycentric mean (BCM) and Tm values calculated by taking the first derivative, as determined by the associated software package.

#### Freeze-thaw stress test

Samples (1 mg/ml) in PBS were subject to five freeze-thaw cycles by placing samples into a minus 80 °C freezer (30 mins) and thawing at room temperature (15 mins). Analytical samples (10 µl) were taken before and afterwards and injected onto a Superdex 200 Increase 5/150 (Cytiva) using an Agilent 1260 Infinity HPLC system. Detection of aggregates was performed using a multi-angle light scattering (MALS) detector (DAWN HELEOS; Wyatt technology) and refractometer (Optilab TRex; Wyatt technology) coupled in-series to the UV detector. Peak analysis was performed using the software, Astra (Wyatt technology).

**Capillary Isoelectric Focussing (cIEF)**

All samples (8 µg) were desalted (< 50 mM NaCl) and added to a cIEF master mix (including pharmalyte and 1.5-3 M urea) in accordance with manufacturer instructions. The final sample was loaded onto a PA800 plus pharmaceutical analysis system with fitted capillary (neutral-coated) and required running reagents (Beckman Coulter). Analysis was performed using associated software.

**CE-SDS**

Samples (20 µg) were desalted (< 50 mM NaCl) and either alkylated (non-reduced) or reduced (with 2-mercaptoethanol) prior to heating samples for 10 mins at 70 °C, in accordance with manufacturer instructions. The final sample was loaded onto a PA800 plus pharmaceutical analysis system with fitted capillary (bare-fused silica) and required running reagents (Beckman Coulter). Analysis was performed using associated software.

**HPLC-SEC and AC-SINS**

Purified antibodies were run on HPLC-SEC (Agilent 1100) using a Superdex 200 Increase column, with 1X PBS running buffer and a flow rate of 0.25 mL/minute to assess the column retention time. The AC-SINS protocol used in the assay is previously described *(41)*. Antibodies were tested at a concentration of 10 µg/mL. Antibodies that showed >20 nm shift were classified as ‘poor’.

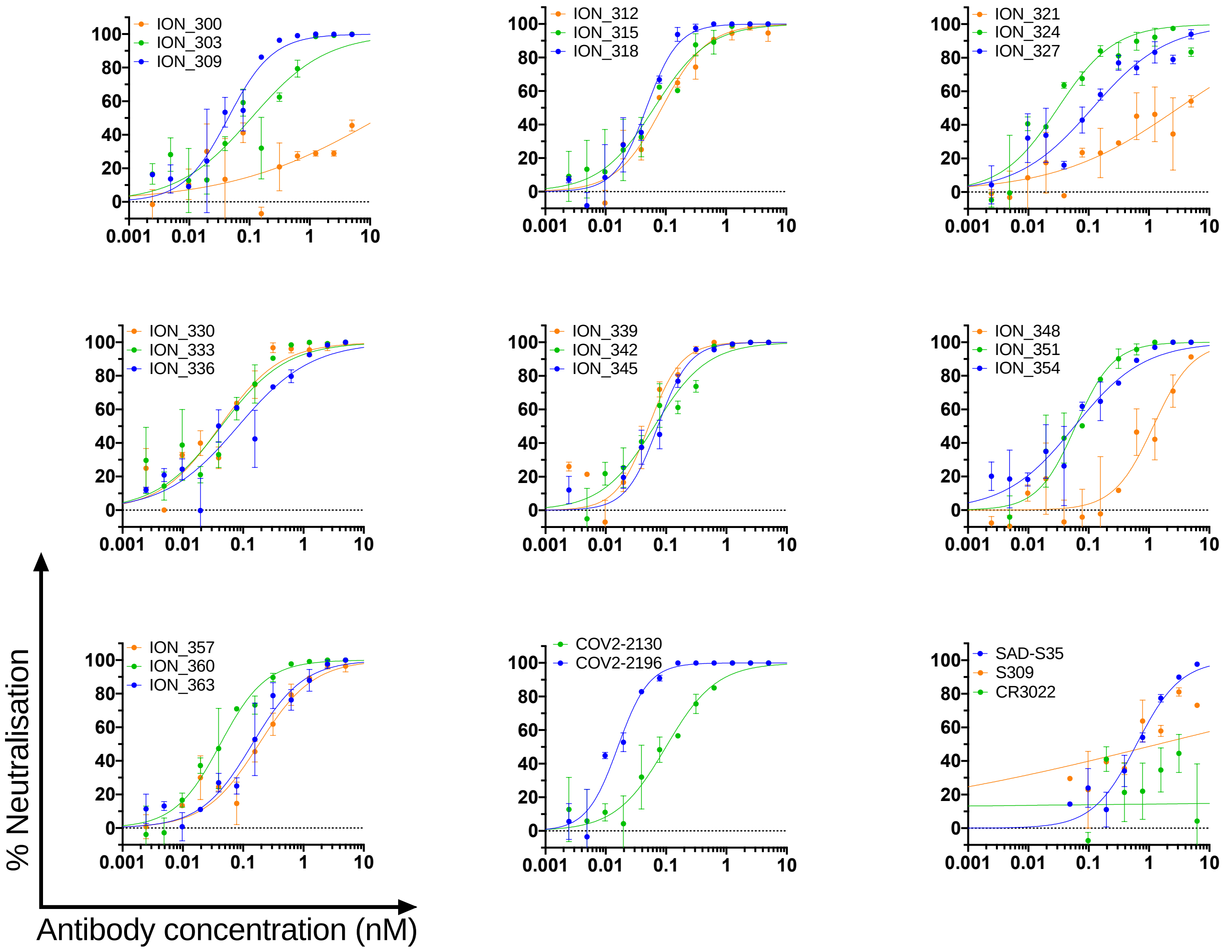


**Fig. S1. Dose-response curves demonstrating pseudovirus neutralisation.** The top panel of 21 antibodies were tested for pseudovirus neutralisation. Control antibodies COV2-2130, COV2-2196, SAD-S35, S309 and CR3022 were included.

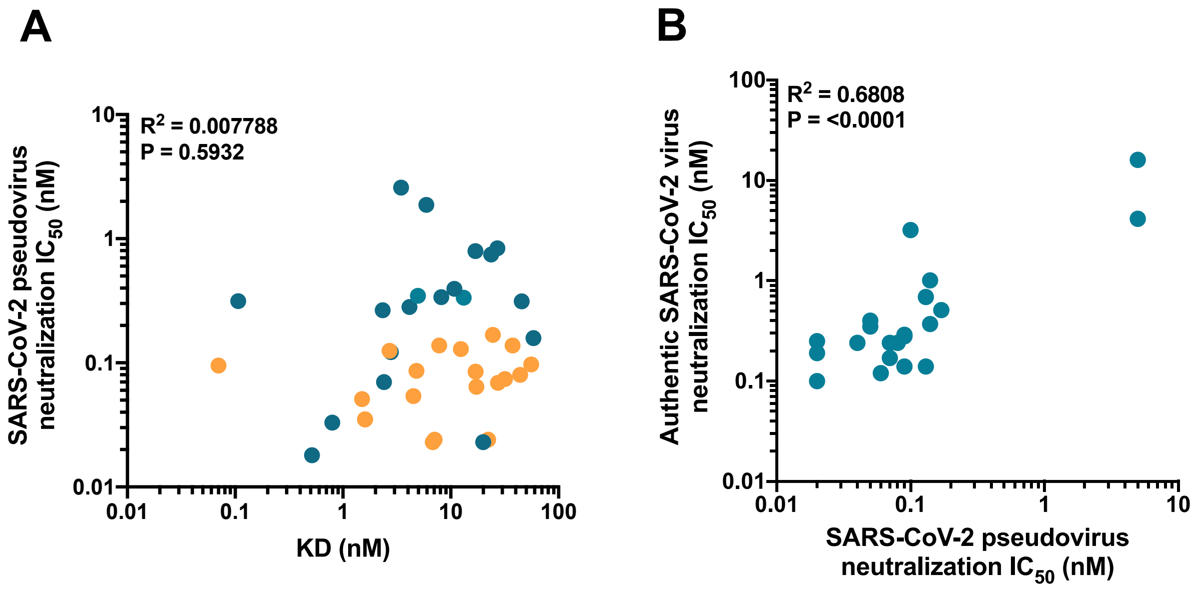


**Fig. S2. Correlation of pseudovirus neutralisation with authentic virus neutralisation and affinity.** A) Correlation between pseudovirus neutralisation and affinity. $IC_{50}$ values plotted (nM) and KD (nM) for a panel of 39 antibodies, 19 of which are in the panel of the top 21 antibodies (shown in orange). B) Correlation between pseudovirus neutralisation and authentic virus neutralisation. $IC_{50}$ values plotted (nM) for each neutralisation assay for the final panel of 21 antibodies. $R^2$ and P values were determined via simple linear regression using GraphPad version 8.

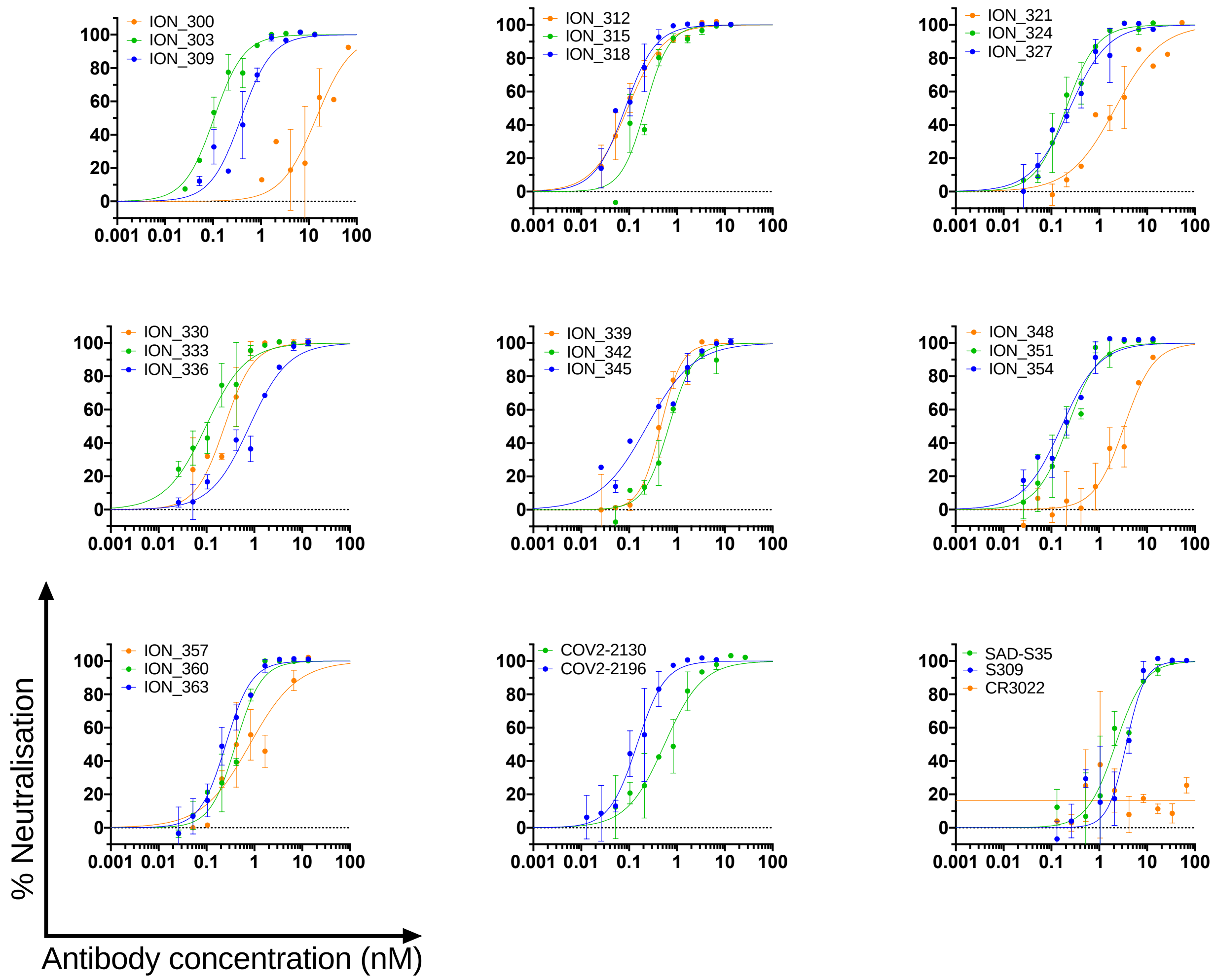


**Fig. S3. Dose-response curves demonstrating authentic virus neutralisation.** The top panel of 21 antibodies were tested for neutralisation of the Australian isolate VIC01/2020 virus strain. Control antibodies COV2-2130, COV2-2196, SAD-S35, S309 and CR3022 were also included.

**A**

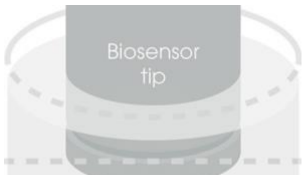

**B**

Antibody 2 (columns) / Antibody 1 (rows)

| Clone | ION_309 | ION_339 | ION_348 | CoV2-2196 | ION_312 | ION_330 | ION_333 | ION_336 | ION_345 | ION_351 | ION_357 | ION_360 | **ION 303** | **ION 315** | **ION 342** | **ION 318** | ION_354 | ION_324 | **ION 327** | **ION 363** | CoV2-2130 | ION_300 | ION_321 |
|---|---|---|---|---|---|---|---|---|---|---|---|---|---|---|---|---|---|---|---|---|---|---|---|
| ION_309 | 0.00 | 0.01 | -0.03 | -0.01 | -0.01 | 0.02 | 0.00 | 0.00 | -0.01 | 0.01 | 0.00 | 0.01 | -0.01 | -0.01 | 0.01 | 0.06 | 0.01 | -0.01 | -0.03 | 0.05 | -0.02 | 0.31 | 0.29 |
| ION_339 | -0.03 | 0.00 | -0.05 | -0.03 | -0.05 | -0.01 | -0.01 | -0.02 | -0.01 | -0.01 | -0.01 | -0.02 | -0.04 | -0.05 | -0.01 | 0.10 | -0.01 | -0.03 | -0.05 | 0.02 | -0.04 | 0.38 | 0.37 |
| ION_348 | 0.00 | 0.00 | -0.04 | -0.01 | -0.01 | 0.01 | 0.01 | 0.00 | -0.01 | 0.01 | -0.01 | 0.02 | 0.00 | -0.01 | 0.02 | 0.08 | 0.01 | -0.01 | -0.02 | 0.08 | -0.01 | 0.13 | 0.11 |
| CoV2-2196 | -0.02 | -0.01 | -0.05 | -0.03 | -0.03 | -0.01 | -0.02 | -0.01 | -0.02 | 0.00 | -0.02 | -0.01 | -0.02 | -0.03 | 0.00 | 0.04 | 0.00 | -0.03 | -0.03 | 0.02 | 0.17 | 0.24 | 0.23 |
| ION_312 | -0.02 | -0.02 | -0.04 | -0.03 | -0.03 | 0.00 | -0.02 | -0.01 | -0.03 | 0.00 | 0.00 | -0.01 | -0.01 | -0.03 | 0.00 | 0.04 | -0.02 | -0.03 | -0.04 | 0.02 | 0.10 | 0.18 | 0.19 |
| ION_330 | -0.02 | -0.01 | -0.03 | -0.02 | -0.04 | 0.00 | -0.01 | -0.02 | -0.02 | -0.01 | 0.00 | -0.01 | -0.04 | -0.04 | 0.00 | 0.04 | 0.02 | -0.02 | -0.03 | 0.02 | 0.19 | 0.29 | 0.27 |
| ION_333 | -0.03 | -0.02 | -0.05 | -0.04 | -0.04 | -0.01 | -0.02 | -0.03 | -0.03 | -0.02 | -0.01 | -0.03 | -0.03 | -0.06 | -0.01 | 0.08 | 0.00 | -0.03 | -0.05 | 0.00 | 0.18 | 0.33 | 0.37 |
| ION_336 | -0.03 | -0.01 | -0.05 | -0.03 | -0.04 | 0.00 | -0.01 | -0.02 | -0.02 | -0.01 | -0.01 | -0.03 | -0.03 | -0.05 | -0.01 | 0.09 | 0.00 | -0.02 | -0.05 | 0.00 | 0.23 | 0.40 | 0.41 |
| ION_345 | -0.03 | -0.02 | -0.06 | -0.04 | -0.04 | 0.00 | -0.01 | -0.03 | -0.03 | -0.02 | -0.01 | -0.03 | -0.03 | -0.03 | -0.02 | 0.03 | -0.02 | -0.03 | -0.06 | -0.02 | 0.11 | 0.22 | 0.22 |
| ION_351 | -0.04 | -0.03 | -0.06 | -0.04 | -0.05 | -0.01 | -0.01 | 0.05 | -0.05 | -0.02 | -0.02 | -0.04 | -0.04 | -0.05 | -0.02 | 0.02 | -0.01 | -0.04 | -0.05 | -0.01 | 0.30 | 0.52 | 0.56 |
| ION_357 | -0.03 | -0.02 | -0.06 | -0.03 | -0.04 | 0.00 | -0.01 | 0.01 | -0.03 | -0.02 | -0.01 | -0.03 | -0.03 | -0.04 | 0.01 | 0.03 | 0.00 | -0.03 | -0.04 | 0.00 | 0.37 | 0.50 | 0.55 |
| ION_360 | -0.04 | -0.03 | -0.06 | -0.04 | -0.03 | -0.01 | -0.01 | 0.02 | -0.03 | -0.02 | -0.02 | -0.04 | -0.03 | -0.04 | -0.02 | 0.01 | -0.02 | -0.04 | -0.06 | -0.02 | 0.21 | 0.35 | 0.34 |
| **ION 303** | -0.01 | -0.01 | -0.03 | 0.00 | -0.01 | 0.01 | 0.00 | -0.01 | -0.02 | 0.01 | 0.00 | 0.01 | 0.00 | -0.02 | 0.01 | 0.05 | 0.01 | -0.02 | -0.02 | 0.03 | 0.00 | 0.18 | 0.19 |
| **ION 315** | -0.02 | -0.02 | -0.06 | -0.04 | -0.03 | -0.01 | 0.00 | 0.01 | -0.03 | -0.01 | -0.01 | -0.02 | -0.02 | -0.03 | -0.01 | 0.03 | -0.02 | -0.04 | -0.05 | 0.00 | 0.07 | 0.16 | 0.18 |
| **ION 342** | -0.03 | -0.01 | -0.05 | -0.03 | -0.04 | 0.00 | -0.01 | -0.02 | -0.03 | -0.01 | -0.01 | -0.02 | -0.02 | -0.03 | -0.01 | 0.04 | -0.02 | -0.03 | -0.05 | -0.02 | 0.08 | 0.17 | 0.21 |
| **ION 318** | -0.02 | -0.02 | -0.03 | -0.04 | -0.02 | 0.01 | -0.01 | -0.01 | -0.02 | 0.00 | 0.00 | -0.02 | 0.00 | -0.03 | -0.01 | 0.02 | -0.03 | -0.02 | -0.03 | 0.00 | 0.00 | 0.31 | 0.31 |
| ION_354 | -0.04 | -0.03 | -0.06 | -0.04 | -0.04 | -0.01 | -0.01 | 0.06 | -0.04 | -0.03 | -0.02 | -0.04 | -0.04 | -0.04 | -0.01 | 0.02 | -0.01 | -0.04 | -0.06 | 0.12 | 0.21 | 0.35 | 0.40 |
| ION_324 | -0.01 | 0.00 | -0.02 | 0.00 | -0.02 | 0.01 | 0.00 | -0.01 | -0.01 | 0.00 | 0.00 | 0.01 | -0.01 | -0.02 | 0.02 | 0.07 | 0.02 | -0.01 | -0.02 | 0.15 | 0.01 | 0.13 | 0.13 |
| **ION 327** | -0.01 | 0.00 | -0.02 | -0.01 | -0.02 | 0.01 | 0.00 | 0.00 | -0.01 | 0.00 | -0.01 | 0.00 | -0.01 | -0.03 | 0.01 | 0.06 | 0.01 | -0.01 | -0.02 | 0.13 | 0.00 | 0.09 | 0.08 |
| **ION 363** | 0.01 | 0.00 | -0.06 | -0.03 | -0.03 | -0.01 | 0.00 | 0.02 | -0.03 | -0.02 | -0.02 | -0.03 | 0.00 | -0.02 | -0.01 | 0.01 | 0.06 | 0.47 | 0.10 | -0.03 | 0.18 | 0.37 | 0.30 |
| CoV2-2130 | -0.01 | -0.02 | -0.04 | 0.25 | 0.26 | 0.28 | 0.26 | 0.22 | 0.22 | 0.26 | 0.18 | 0.23 | 0.12 | 0.26 | 0.26 | 0.18 | 0.25 | -0.02 | -0.03 | 0.24 | -0.01 | 0.24 | 0.25 |
| ION_300 | 0.62 | 0.59 | 0.53 | 0.57 | 0.64 | 0.66 | 0.60 | 0.45 | 0.50 | 0.57 | 0.40 | 0.51 | 0.71 | 0.64 | 0.62 | 0.70 | 0.53 | 0.63 | 0.62 | 0.53 | 0.29 | -0.01 | -0.01 |
| ION_321 | 0.44 | 0.48 | 0.38 | 0.51 | 0.57 | 0.61 | 0.51 | 0.38 | 0.41 | 0.51 | 0.35 | 0.45 | 0.61 | 0.58 | 0.53 | 0.61 | 0.41 | 0.55 | 0.19 | 0.41 | 0.15 | -0.03 | -0.03 |

**Fig. S4. Epitope binning with RBD on panel of 21 antibodies and controls.** A) Assay format. Using Octet BLI technology, in both orientations (i.e. Capture vs Detector) a classical sandwich assay format was employed to identify the presence of seven different epitope bins. Two characterized control antibodies (COV2-2130 and COV2-2196) were also included in the analyses. B) Epitope binning results matrix. All antibodies were tested as Ab1 – Capture and as Ab2 – Detector. The signals are organised in a matrix, where the self-self pairing responses are displayed along the diagonal. Responses above 0.1nm were considered as positive (i.e. presence of viable pairing).

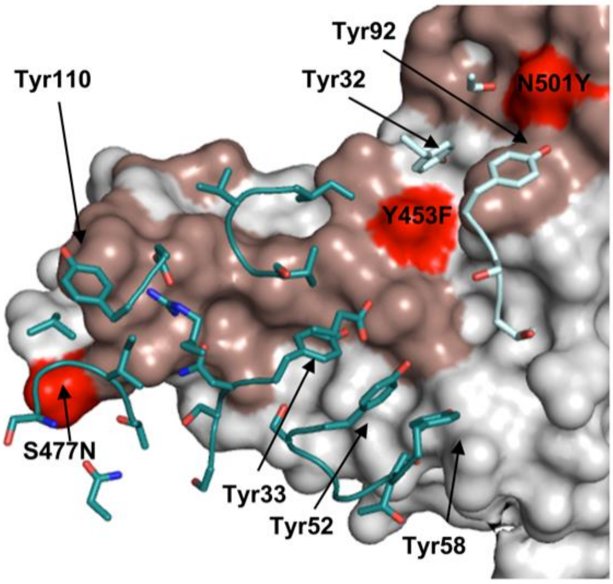


**Fig. S5. Analysis of the ION_360:RBD Interface.** CDR residues within 5 Å of the RBD are shown in sticks for the $V_H$ (teal) and $V_K$ (cyan) chains. The RBD is represented as a surface (grey) with ACE2 binding residues highlighted (salmon). Known SARS-CoV-2 mutations within the receptor binding motif (Y453F, S477N and N501Y), are highlighted (red). The RBD:ION_360 interface buries 851 Å$^2$ of RBD protein surface from bulk solvent following complex formation. Of the 20 RBD residues within 5 Å of the ACE2 receptor that make up the binding site, 13 are buried upon ION_360 binding. In contrast, 859 Å$^2$ of antibody surface is buried from bulk solvent, with 712 Å$^2$ of the buried surface from the $V_H$ domain and 147 Å$^2$ from the $V_K$ domain. With the exception of CDR-L2, at least one residue from all CDR loops is within 5 Å of the SARS-CoV-2 RBD, with 22 amino acids from the CDR loops losing at least 10 Å$^2$ of solvent accessibility and making direct contacts with the protein. The interface features a mixture of polar and hydrophobic contacts, including the involvement of six aromatic side chains in the CDR loops, from both the $V_H$ (Tyr33, Tyr52, Tyr58, Tyr110) and $V_K$ (Tyr32 and Tyr92) domains, as well as a network of 13 hydrogen bonds, accounting for a measured $K_D$ of 1.5 nM.

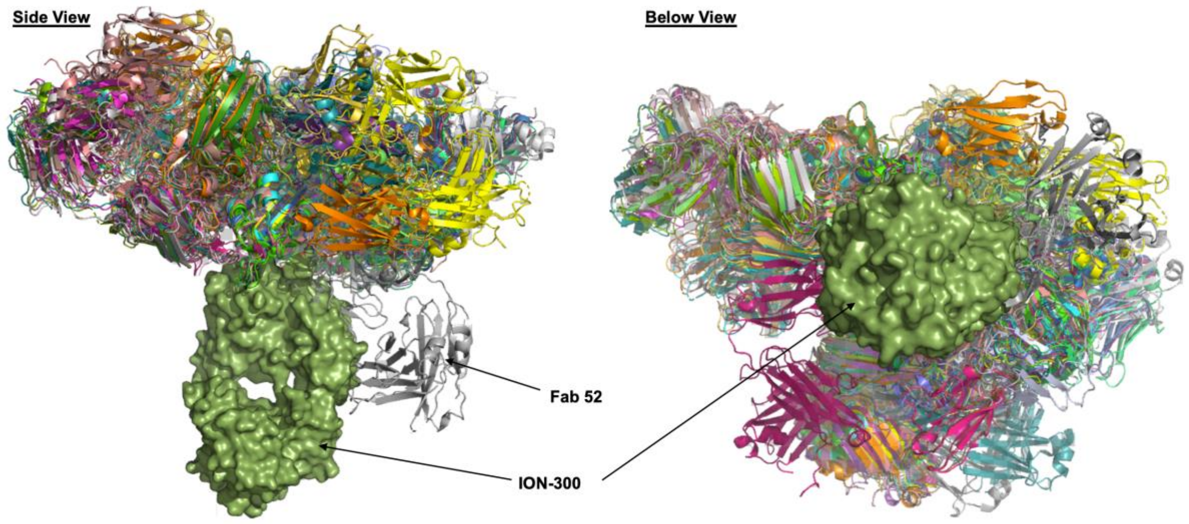


**Fig. S6. Superposition of antibody and nanobody structures targeting the SARS-CoV-2 RBD.** Cartoon representation of all published antibody and nanobody structures that bind to the SARS-CoV-2 RBD, superposed onto the RBD of the ION_300: RBD structure (green molecular surface), from the side and below viewpoints of the ION_300 molecule.

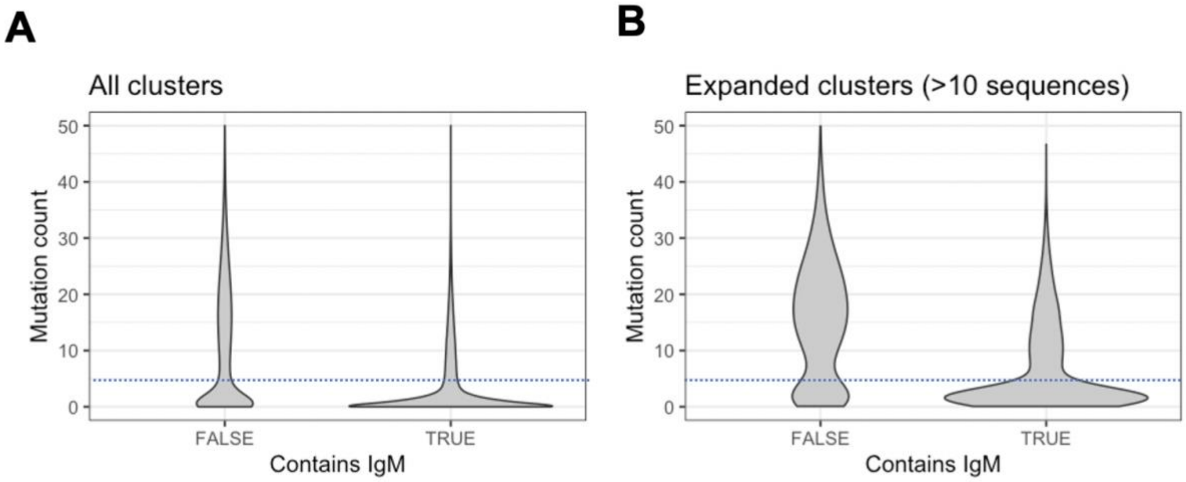


**Fig. S7. Mutation distribution of clusters.** A) Clusters were stratified according to whether they contained any IgM sequences. Violin plot shows the average mutation of the clusters within each group. B) Same as A, but only including clusters that contain >10 sequences. For A and B, the dotted line indicates the cutoff that was used to distinguish naïve, or recently activated cells from memory cells.

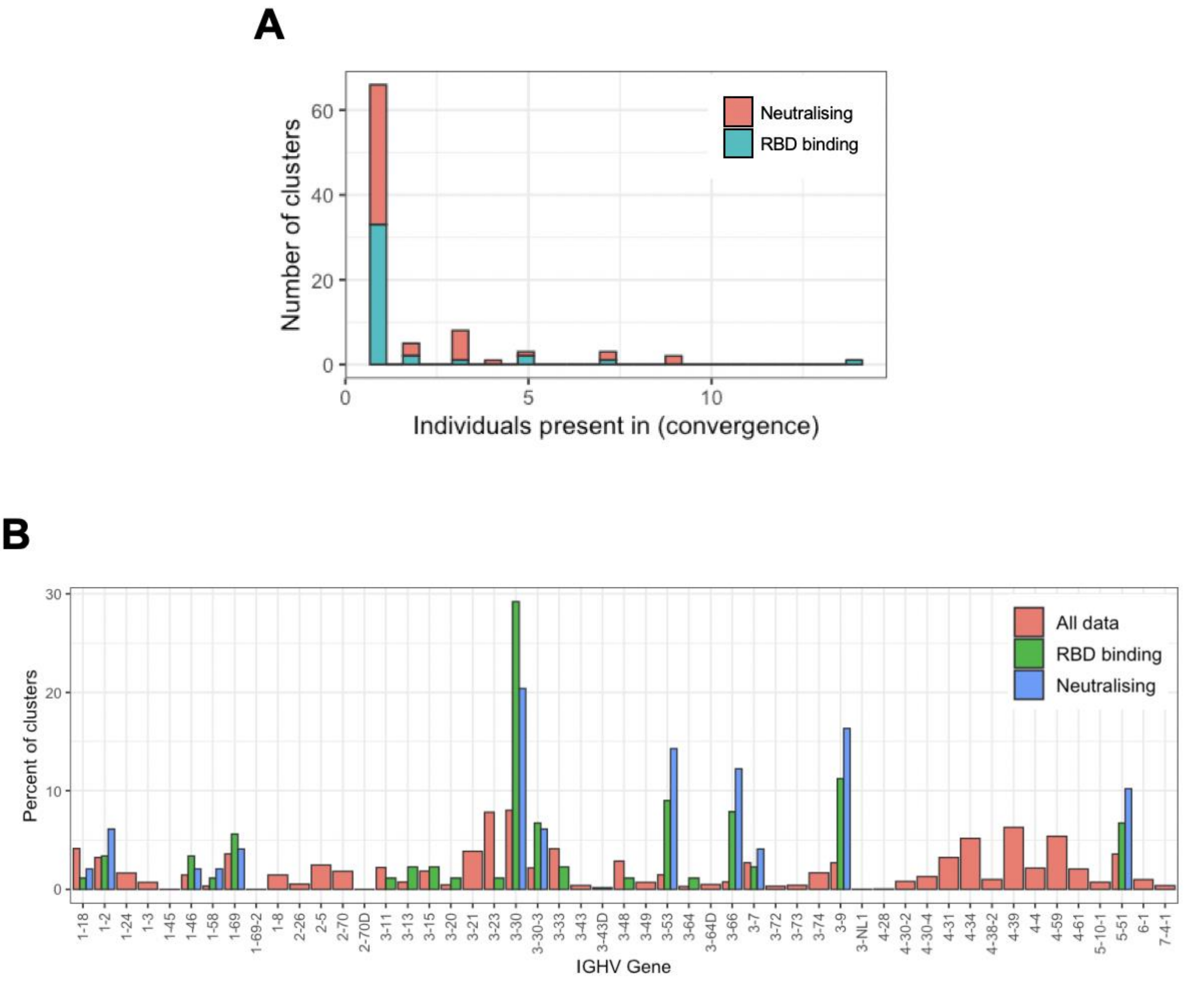


**Fig. S8. VH germline usage in convergent antibody response**. A) The number of individuals the clusters annotated as RBD-binding or RBD binding and neutralising are present in. B) V gene segment usage distribution of clusters.

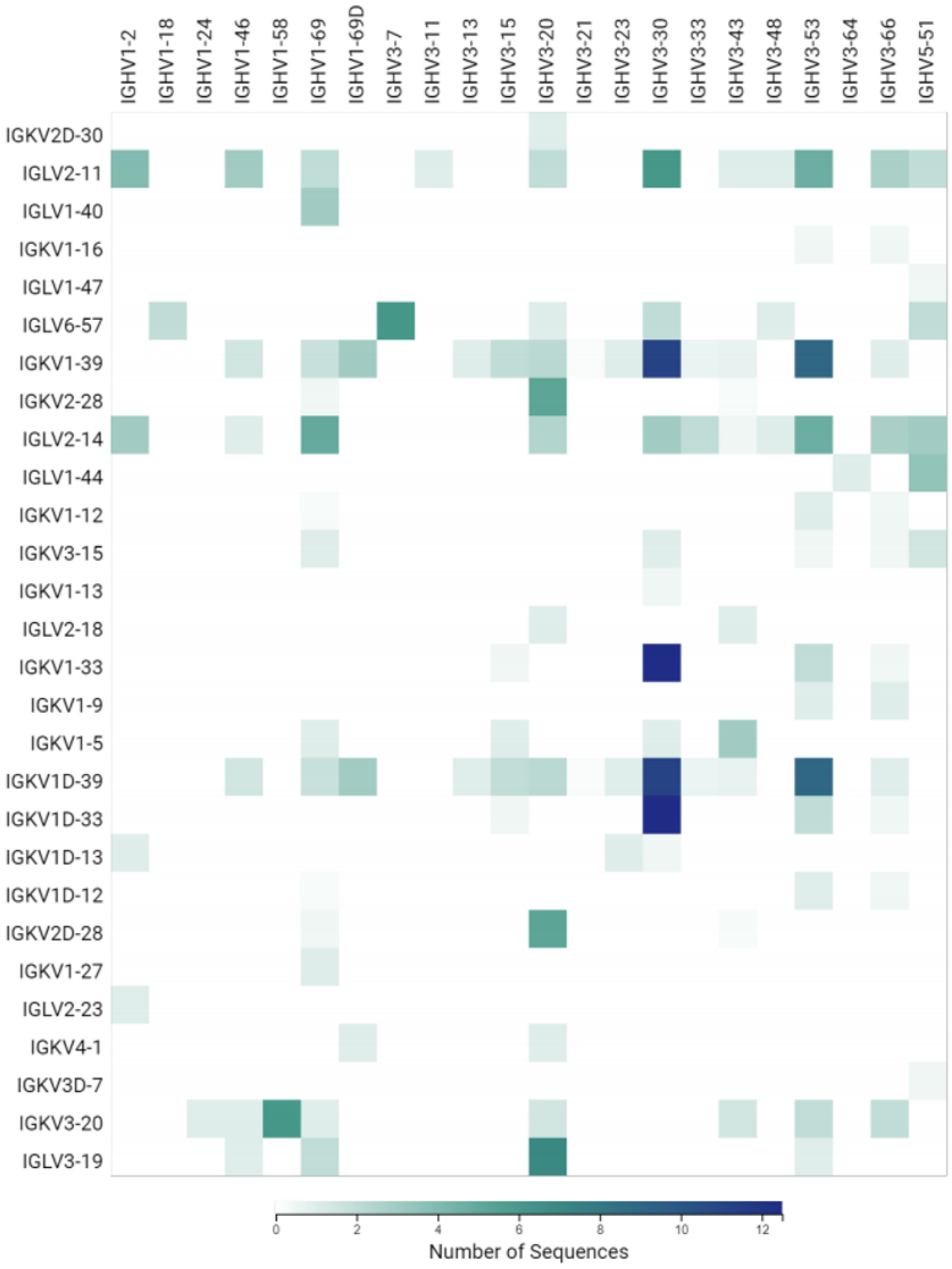


**Fig. S9. Heavy and light chain V gene utilisation and pairing preference of all RBD-binding antibodies isolated using phage display technology.** Analysis performed with Geneious Biologics (Biomatters).

| Reference | No. of days of symptoms (mean) prior to sample collection |
|---|---|
| Rogers et al (2020) | 17 |
| Hansen et al (2020) | 24.5 |
| Zost et al[A] (2020) | 50 |
| Cao et al (2020) | 38 |
| Zost et al[B] (2020) | 43 |
| Liu et al (2020) | 25 |
| Kreer et al (2020) | 25 |
| M. Brouwer et al (2020) | 28 |
| Bin Ju et al (2020) | 17 |
| Robbiani et al (2020) | 39 |
| Tortorici et al (2020) | 53.5 |
| Mean | 32.7 |

**Table S1. Days of symptoms prior to sample collection in related studies.**

| Donor ID | Age group | Gender | Ethnicity | No. of days of symptoms prior to sample collection | Clinical status |
|---|---|---|---|---|---|
| BARTS_01 | 30-39 | Female | Caucasian | 7 | Deteriorating |
| BARTS_02 | 30-39 | Male | SE Asian | 10 | Stable |
| BARTS_03 | 50-59 | Male | Caucasian | 11 | Stable |
| BARTS_04 | 70-79 | Male | Black | 14 | Stable |
| BARTS_05 | 70-79 | Female | SE Asian | 7 | Stable |
| BARTS_06 | 30-39 | Male | N/A | 14 | Improving |
| BARTS_07 | 30-39 | Male | N/A | 4 | Improving |
| BARTS_08 | 50-59 | Male | SE Asian | 15 | Stable |
| BARTS_10 | 70-79 | Female | Caucasian | 10 | Stable |
| BARTS_11 | 20-29 | Male | SE Asian | 9 | Stable |
| BARTS_12 | 80-89 | Female | Caucasian | Uncertain | Stable |
| BARTS_13 | 20-29 | Male | Asian | 11 | Stable |
| BARTS_14 | 40-49 | Male | Caucasian | 6 | Deteriorating |
| BARTS_15 | 30-39 | Male | SE Asian | 11 | Stable |
| BARTS_16 | 50-59 | Female | N/A | 11 | Stable |
| BARTS_17 | 40-49 | Female | Black | 12 | Stable |
| BARTS_18 | 30-39 | Male | Caucasian | 20 | Improving |
| BARTS_19 | 40-49 | Male | SE Asian | 15 | Improving |

**Table S2. Clinical details and characteristics of the patient cohort used in this study**. In this cohort, 12 (66.7%) patients were male and 6 were females (33.3%), and had a mean age of 48.8 (range 25.6-87.4) years. The clinical status of the patient on the day of sample collection was subjectively assigned as Improving, Stable or Deteriorating by the direct clinical care team, on the basis of increasing, stable, or decreasing requirement of supplemental oxygen in comparison to the previous three days.

| Antibody ID | $K_{on}$ ($M^{-1}s^{-1}$) | $K_{off}$ ($s^{-1}$) | KD (nM) |
|---|---|---|---|

| | | | |
|---|---|---|---|
| ION_300 | $2.52 \times 10^5$ | $7.75 \times 10^{-4}$ | 3.4 |
| ION_303 | $9.93 \times 10^5$ | $4.60 \times 10^{-3}$ | 4.8 |
| ION_309 | $5.57 \times 10^5$ | $4.27 \times 10^{-3}$ | 7.8 |
| ION_312 | $1.18 \times 10^6$ | $1.44 \times 10^{-2}$ | 12.4 |
| ION_315 | $6.73 \times 10^5$ | $1.14 \times 10^{-2}$ | 17.0 |
| ION_318 | $6.97 \times 10^5$ | $4.70 \times 10^{-3}$ | 6.8 |
| ION_321 | $4.26 \times 10^5$ | $2.73 \times 10^{-4}$ | 0.7 |
| ION_324 | $1.00 \times 10^6$ | $2.99 \times 10^{-2}$ | 31.9 |
| ION_327 | $9.86 \times 10^5$ | $4.20 \times 10^{-2}$ | 44.0 |
| ION_330 | $3.27 \times 10^5$ | $8.82 \times 10^{-3}$ | 27.4 |
| ION_333 | $2.38 \times 10^5$ | $4.11 \times 10^{-3}$ | 17.3 |
| ION_336 | $8.03 \times 10^4$ | $1.22 \times 10^{-3}$ | 37.5 |
| ION_339 | $3.85 \times 10^5$ | $1.71 \times 10^{-3}$ | 4.5 |
| ION_342 | $3.12 \times 10^5$ | $7.65 \times 10^{-3}$ | 24.6 |
| ION_345 | $1.76 \times 10^5$ | $3.71 \times 10^{-3}$ | 22.2 |
| ION_348 | $5.79 \times 10^5$ | $3.12 \times 10^{-2}$ | 55.7 |
| ION_351 | $1.86 \times 10^5$ | $2.91 \times 10^{-4}$ | 1.6 |
| ION_354 | $1.73 \times 10^5$ | $1.23 \times 10^{-3}$ | 7.1 |
| ION_357 | $1.09 \times 10^5$ | $2.42 \times 10^{-4}$ | 2.7 |
| ION_360 | $1.88 \times 10^5$ | $2.79 \times 10^{-4}$ | 1.5 |
| ION_363 | $6.12 \times 10^5$ | $4.39 \times 10^{-5}$ | 0.07 |

**Table S3. 1:1: binding kinetics of the final panel of 21 antibodies to SARS-CoV-2 RBD measured by SPR.**

| Antibody ID | Pseudovirus neutralisation $IC_{50}$ (nM) | Authentic virus neutralisation $IC_{50}$ (nM) |
|---|---|---|
| ION_300 | >5 | 16.00 |
| ION_303 | 0.09 | 0.14 |
| ION_309 | 0.14 | 0.37 |
| ION_312 | 0.13 | 0.14 |
| ION_315 | 0.09 | 0.28 |
| ION_318 | 0.02 | 0.10 |
| ION_321 | >5 | 4.15 |
| ION_324 | 0.07 | 0.24 |
| ION_327 | 0.08 | 0.24 |
| ION_330 | 0.07 | 0.17 |
| ION_333 | 0.06 | 0.12 |
| ION_336 | 0.14 | 1.01 |
| ION_339 | 0.05 | 0.40 |
| ION_342 | 0.17 | 0.51 |
| ION_345 | 0.02 | 0.25 |
| ION_348 | 0.10 | 3.20 |
| ION_351 | 0.04 | 0.24 |
| ION_354 | 0.02 | 0.19 |
| ION_357 | 0.13 | 0.69 |
| ION_360 | 0.05 | 0.35 |
| ION_363 | 0.09 | 0.29 |
| COV2-2130 | 0.10 | 0.84 |
| COV2-2196 | 0.02 | 0.12 |
| S309 | 1.42 | 2.40 |

**Table S4. Neutralisation potency of the selected 21 antibodies against a pseudotyped virus and an authentic strain of SARS-CoV-2.** Neutralising antibody titre is expressed as 50% inhibitory concentration ($IC_{50}$). The strain of authentic virus used was SARS-CoV-2 Australia/VIC01/2020. Control antibodies COV2-2130, COV-2196 and S309 were also included.

| Antibody ID | Freeze-thaw stress | pH stress | Thermal stress | Capillary isoelectric focusing (cIEF) | | Non-reduced CE-SDS | | HPLC-SEC profile | AC-SINS shift (nm) | Overall ranking |
|---|---|---|---|---|---|---|---|---|---|---|
| | Loss (%) | Loss after Protein A (%) | Tm (°C) | Main group (pI) | | Glycosylated intact antibody (%) | Multiple species | | | |
| | | | | Lower | Upper | | | | | |
| ION_300 | 0 | 4.3 | 63.2 | 8.46 | 8.63 | 97.6 | No | S | 7 | PASS |
| ION_303 | 0.8 | 4.2 | 68.9 | 6.84 | 7.02 | 93.4 | No | S | 5 | FAIL |
| ION_309 | 0.1 | 2.5 | 73.9 | 8.58 | 8.66 | 91.9 | No | S | 11 | PASS |
| ION_312 | 0.8 | 9.3 | 70.6 | 8.31 | 8.41 | 90.1 | No | S | 4 | PASS |
| ION_315 | 1.3 | 4.8 | 68.5 | 7.55 | 7.66 | 94.4 | No | S | 4 | PASS |
| ION_318 | 0 | 1.9 | 71.3 | 8.77 | 8.83 | 87.2 | No | D | 29 | FAIL |
| ION_321 | 0 | 3.6 | 64.1 | 8.45 | 8.60 | 68.6 | Yes | S | 7 | FAIL |
| ION_324 | 0 | 1 | 64.7 | 7.72 | 7.93 | 67.6 | Yes | S | 4 | FAIL |
| ION_327 | 0 | 5.4 | 67 | 7.45 | 7.58 | 60.4 | Yes | D | 7 | FAIL |
| ION_330 | 0 | 2.4 | 62.8 | 7.72 | 7.82 | 96.5 | No | S | 6 | PASS |
| ION_333 | 0.3 | 1.2 | 74.5 | 7.32 | 7.38 | 86.1 | No | S | 4 | FAIL |
| ION_336 | 0.3 | 3.9 | 74.2 | 8.47 | 8.57 | 93.9 | No | S | 4 | PASS |
| ION_339 | 0.9 | 6.2 | 67.9 | 7.63 | 7.83 | 95.1 | No | S | 17 | PASS |
| ION_342 | 0.5 | 3.4 | 72.3 | 8.96 | 9.08 | 94.9 | No | S | 5 | PASS |
| ION_345 | 1 | 2.6 | 68.6 | 8.37 | 8.49 | 91.4 | No | S | 4 | PASS |
| ION_348 | 0.3 | 11.3 | 69 | 7.55 | 7.65 | 95.0 | No | S | 4 | FAIL |
| ION_351 | 0 | 2.4 | 71.1 | 8.58 | 8.74 | 93.1 | No | S | 4 | PASS |
| ION_354 | 0.3 | 3 | 77.1 | 8.93 | 9.08 | 94.0 | No | S | 5 | PASS |
| ION_357 | 0.5 | 2.4 | 68.6 | 9.00 | 9.11 | 90.8 | No | S | 4 | PASS |
| ION_360 | 0.4 | 4.7 | 65.1 | 8.85 | 9.01 | 90.2 | No | S | 7 | PASS |
| ION_363 | 0 | 5.3 | 63.3 | 6.83 | 6.91 | 67.7 | Yes | S | 9 | FAIL |

**Table S5. Summary of developability data for final panel of 21 antibodies**. Cut-offs were given for each test to decide what defines a "passing" antibody: <5% freeze-thaw loss, <10% loss after protein A, Tm >60°C, lower (<7.5) and upper pI species (>9.5), >90% intact glycosylated antibody and no presence of multiple species in CE-SDS, a standard HPLC-SEC profile, AC-SINS shift <20 nm. HPLC-SEC profiles defined by retention time (Early ('E') – 1.47 minutes or earlier, Standard ('S') – 1.48-1.56 minutes, Delayed ('D')– 1.57 minutes or more). Antibodies that passed criteria set were given a green fill colour; antibodies that failed the criteria set were given an orange colour. A yellow fill colour was given where antibodies were considered weaker but were not classified as failed. An overall developability pass or fail ranking was given based on all criteria.

| | RBD:ION_300 | RBD:ION_360 |
|---|---|---|
| ***Data Collection*** | | |
| Beamline | ID30A1 | ID30A1 |
| Wavelength (Å) | 0.96861 | 0.97950 |
| Space Group | $P4_3$ | $P2_1$ |
| Cell Dimensions<br>a, b, c (Å), α, β, γ (°) | 75, 75, 143, 90, 90, 90 | 91, 108, 182, 90, 99, 90 |
| Resolution (Å) | 47.70 – 2.35 | 48.28 – 2.80 |
| $R_{merge}$ | 0.051 (0.850) | 0.074 (0.553) |
| $CC_{1/2}$ | 0.993 (0.489) | 0.990 (0.491) |
| I/σI | 17.5 (1.5) | 5.6 (1.3) |
| Completeness (%) | 99.1 (99.9) | 99.9 (99.1) |
| Redundancy | 3.5 (3.5) | 2.0 (2.0) |
| ***Refinement*** | | |
| No. of Reflections | 114774 (11217) | 170770 (17020) |
| No. of Unique | 32467 (3234) | 85556 (8446) |
| $R_{factor}$ / $R_{free}$ (%) | 21.5 (26.2) | 24.1 (29.1) |
| Wilson *B*-factors (Å) | 56.5 | 67.9 |
| *B*-factors (Å) | | |
| Protein | 72.5 | 69.4 |
| R.M.S. Deviations | | |
| Bond lengths (Å) | 0.005 | 0.011 |
| Bond angles (°) | 0.860 | 1.680 |
| Ramachandran Favoured (%) | 95.84 | 92.7 |

**Table S6. X-ray data and refinement statistics.**
